# Structural basis for receptor selectivity and inverse agonism in S1P_5_ receptors

**DOI:** 10.1101/2022.02.25.480536

**Authors:** Elizaveta Lyapina, Egor Marin, Anastasiia Gusach, Philipp Orekhov, Andrey Gerasimov, Aleksandra Luginina, Daniil Vakhrameev, Margarita Ergasheva, Margarita Kovaleva, Georgii Khusainov, Polina Khorn, Mikhail Shevtsov, Kirill Kovalev, Ivan Okhrimenko, Petr Popov, Hao Hu, Uwe Weierstall, Wei Liu, Yunje Cho, Ivan Gushchin, Andrey Rogachev, Gleb Bourenkov, Sehan Park, Gisu Park, Hyo Jung Hyun, Jaehyun Park, Valentin Gordeliy, Valentin Borshchevskiy, Alexey Mishin, Vadim Cherezov

## Abstract

The bioactive lysophospholipid sphingosine-1-phosphate (S1P) acts via five different subtypes of S1P receptors (S1PR) - S1P_1-5_. S1P_5_ is predominantly expressed in nervous and immune systems, regulating the egress of natural killer cells from lymph nodes and playing a role in immune and neurodegenerative disorders, as well as carcinogenesis. Several S1PR therapeutic drugs have been developed to treat these diseases; however, they lack receptor subtype selectivity, which leads to side effects. In this article, we describe a 2.2 Å resolution room temperature crystal structure of the human S1P_5_ receptor in complex with a selective inverse agonist determined by serial femtosecond crystallography (SFX) at the Pohang Accelerator Laboratory X-Ray Free Electron Laser (PAL-XFEL) and analyze its structure-activity relationship data. The structure demonstrates a unique ligand-binding mode, involving an allosteric subpocket, which clarifies the receptor subtype selectivity and provides a template for structure-based drug design. Together with previously published S1PR structures in complex with antagonists and agonists, the new S1P_5_-inverse agonist structure sheds light on the activation mechanism and reveals structural determinants of the inverse agonism in the S1PR-family.

## INTRODUCTION

Sphingosine-1-phosphate (S1P) is a lysosphingolipid bio-regulator produced from ceramide in activated platelets, injured cells, and cells stimulated by protein growth factors^1,2^. S1P is released in the blood^3^, where it regulates angiogenesis^4^, cell proliferation, migration, and mitosis^5^ by activating five subtypes of the S1P G protein-coupled receptors - S1P_1-5_. S1P_1_ couples only to G_i_ protein, S1P_4_ and S1P_5_ signal through G_i_ and G_12/13_^6^, and both S1P_2_ and S1P_3_ couple to G_i_, G_12/13_, and G_q_^7^. S1P receptors have different expression profiles – S1P_1_-S1P_3_ are expressed in all organs throughout the body, while S1P_4_ expression is limited to the immune system, and S1P_5_ is predominantly expressed in the nervous (oligodendrocytes) and immune (NK cells) systems^8^. S1P_5_ also inhibits PAR-1 mediated platelet activation^9^. This receptor plays an important role in autoimmune^10^ and neurodegenerative disorders^10,11^ as well as carcinogenesis^12^. For example, S1P_5_ agonists elicit neuroprotective effects in Alzheimer’s and Huntington’s diseases^10^, while S1P_5_ inhibition leads to apoptosis of cancerous NK cells in large granular leukemia (LGL)^12^. Non-selective modulators such as fingolimod^13^, as well as dual S1P_1_/S1P_5_ ligands siponimod^14^ and ozanimod^15,16^, have been approved for the treatment of multiple sclerosis^17^, Crohn’s disease^18^, and other autoimmune disorders. However, the exact pharmacological role of S1P_5_ remains unclear, mostly due to the lack of well-characterized potent and highly selective S1P_5_ ligands with *in vivo* activity. While inhibition of S1P_5_ is considered as a prospective treatment for LGL^12^, antagonism of S1P_1_ leads to serious adverse effects such as lung capillary leakage, renal reperfusion injury, and cancer angiogenesis^19^. Therefore, high-resolution structures of S1P_5_ in complex with highly selective ligands would shed light on receptor selectivity and provide templates for structure-based design of selective therapeutic drugs with more focused function and fewer side effects.

The first crystal structure of an S1P receptor was published in 2012^20^, revealing the inactive state conformation of the human S1P_1_ in complex with a selective antagonist sphingolipid mimetic ML056. Recently, a crystal structure of S1P_3_ bound to its endogenous agonist^21^, as well as cryo-EM structures of S1P_1_, S1P_3_, and S1P_5_ in complex with G_i_ proteins^22,23^, and S1P_1_ in complex with G_i_ and β-arrestin^24^, provided insights in the activation mechanism for the S1P receptor family. However, no structures of this family members in complex with inverse agonists have been reported to date.

In this article, we present the crystal structure of S1P_5_ in complex with a selective inverse agonist ONO-5430608^25^ determined by serial femtosecond crystallography (SFX) and analyze it alongside structureactivity relationship data from cell-based functional assays using extensive mutagenesis, molecular docking, molecular dynamics, and AlphaFold simulations.

## RESULTS

### Structure determination using an X-ray free-electron laser (XFEL)

Human S1P_5_ receptor was engineered for crystallization by fusing a thermostabilized apocytochrome b562RIL^26^ into the third intracellular loop (ICL3) and adding a haemagglutinin signal peptide, FLAG tag, and a linker on the N terminus as well as a PreScission Protease site and decahistidine tag on the C terminus (Supplementary Fig. 1). Crystals of S1P_5_ bound to an inverse agonist ONO-5430608 were obtained by lipidic cubic phase (LCP) crystallization^27^ reaching a maximum size of 30 μm. Attempts to solve the structure were first made using cryocooled crystals at a synchrotron source achieving a maximum of 4 Å single-crystal data resolution, however, the obtained data could not be phased using molecular replacement. Crystals of S1P_5_ bound to ONO-5430608 were then optimized to grow at a high crystal density with an average size of ~5-10 μm and used for room temperature SFX data collection at PAL-XFEL (Supplementary Fig. 2). The crystal structure was solved at a 2.2 Å resolution in the P212121 space group with identical unit cell constants as the single-crystal data (Supplementary Table 1). A high systematic background scattering from the direct XFEL beam (Supplementary Fig. 3) combined with pseudotranslation led to high structure refinement R-factors, although it did not affect the excellent quality of electron density maps (see Methods and Supplementary Fig. 4). The receptor crystallized with two monomers per asymmetric unit, forming an antiparallel dimer through the TM4-TM4 interface (Supplementary Fig. 5).

### Inactive conformation of S1P_5_ in complex with ONO-5430608

#### Overall architecture

The S1P_5_ structure in complex with ONO-5430608 shares the classical architecture with other class A α-branch lipid receptors^20,21,28^, including a heptahelical transmembrane bundle (7TM), two pairs of disulfide bonds stabilizing extracellular loops 2 and 3 (ECL2 and ECL3), an amphipathic C-terminal helix 8 running parallel to the membrane on the intracellular side, and an N-terminal helix capping the ligandbinding site. As expected, the receptor is captured in the inactive conformation (Fig. 1a,b) based on its overall alignment with the inactive state S1P_1_ (PDB ID 3V2Y, Cα RMSD = 0.84/0.78 Å on 90% of residues for chains A/B of our S1P_5_ structure) and the active state S1P_5_ (PDB ID 7EW1, Cα RMSD = 1.40/1.40 Å on 90% of residues for chains A/B of our S1P_5_ structure) as well as on the conformation of conserved activation-related motifs described in the following section.

**Fig. 1.**
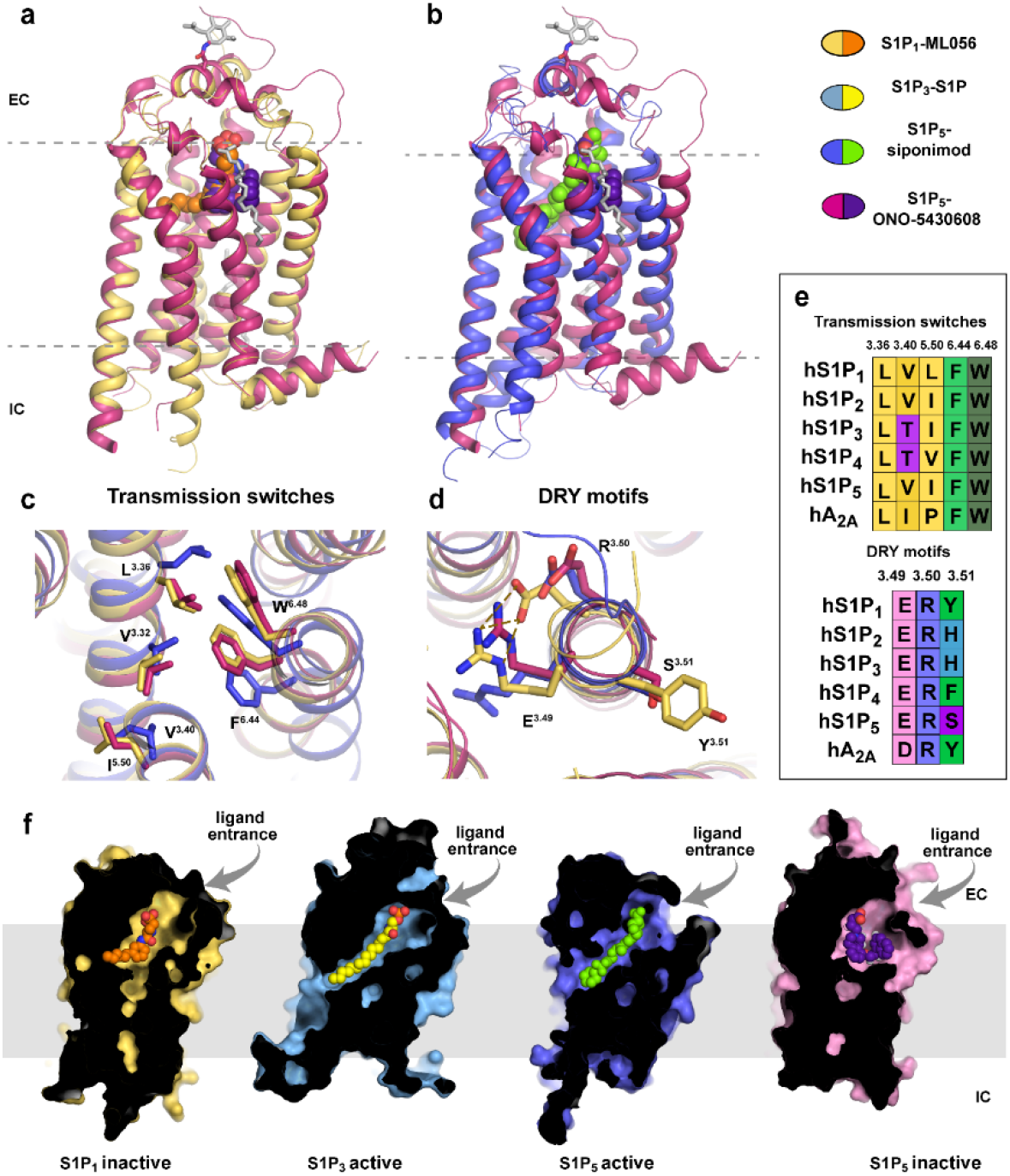
Structure of S1P_5_ and its comparison with structures of other S1PRs: overview and conservative motifs. **a** Superposition of the obtained in this work inactive S1P_5_ structure (pink cartoon) in complex with ONO-5430608 (purple spheres) with the inactive S1P_1_ (yellow cartoon)-ML056 (orange spheres) complex (PDB ID 3V2Y). **b** Superposition of the inactive S1P_5_-ONO-5430608 with the active S1P_5_ (violet cartoon)-siponimod (green spheres) complex (PDB ID 7EW1). Glycosylated residues and lipids observed in the S1P_5_-ONO-5430608 structure are shown as gray sticks. **c** Superposition of the dual toggle switch and P-I-F motif for S1P_5_-ONO-5430608 (inactive state), S1P_5_-siponimod (active state), and S1P_1_-ML056 (inactive state). **d** Superposition of the DRY functional motif for the same three receptorligand pairs as in (**c**). **e** Sequence conservation of the transmission switches and DRY motif in the SP1R family. Adenosine A2A receptor is included as a representative receptor of class A GPCR. **f** Sliced surface representation of known structures from the S1PR family with corresponding ligands: S1P_1_-ML056 (PDB ID 3V2Y), S1P_3_-S1P (PDB ID 7C4S), S1P_5_-siponimod (PDB ID 7EW1), and S1P_5_-ONO5430608 (this work, PDB ID 7YXA).

#### Activation-related conserved motifs and Na-binding site

A dual toggle switch L(F)^3.36^-W^6.48^ (superscripts refer to the Ballesteros-Weinstein^29^ residue numbering scheme in class A GPCRs) together with P^5.50^-I^3.40^-F^6.44^ motif have been characterized as the common microswitches in class A GPCRs that transmit activation-related conformational changes from the ligandbinding pocket towards an outward movement of TMs 5 and 6 and inward displacement of TM7 on the intracellular side^30,31^. In all S1P receptors, the dual toggle switch is conserved as L^3.36^-W^6.48^; however, the first two residues of the P-I-F motif deviate from the consensus (Fig. 1e). Nevertheless, the I^5.50^-V^3.40^-F^6.44^ motif in S1P_5_ appears to serve a similar role as the classical P-I-F motif in other receptors, as the sidechains of V^3.40^ and F^6.44^ switch over upon activation. The I-V-F switch in S1P_5_ is connected to the dual toggle switch through steric interactions between F^6.44^ and W^6.48^, and the shift of W264^6.48^ is accompanied by a rotamer switch of L119^3.36^ (Fig. 1c). Similar dual (also known as “twin”) toggle switch L(F)^3.36^-W^6.48^ has been shown to play a key role in the activation of several other receptors, such as CB1^32,33^, AT1^34^, and MC4^35^.

An allosteric sodium-binding site located in the middle of the 7TM bundle near D^2.50^ is highly conserved in class A GPCRs^36^. Binding of a Na^+^ ion along with several water molecules in this site stabilizes the inactive receptor conformation. Upon receptor activation, the pocket collapses, likely expelling Na^+^ into the cytoplasm^36,37^. Despite a relatively high resolution and conservation of critical sodium binding residues, such as D82^2.50^, S122^3.39^, and N298^7.45^, we could not locate a Na^+^ in the electron density of S1P_5_, most likely because of a low sodium concentration in the final crystallization buffer (~20 mM). Other residues lining the Na^+^-binding pocket (N^1.50^, S^3.39^, N^7.45^, S^7.46^, N^7.49^, Y^7.53^) are also conserved in S1P_5_, with the exception of two polar residues, T79^2.47^ and S81^2.49^, in the side part of the pocket, which are typically represented by two hydrophobic alanines^36^.

On the intracellular side of the receptor, conserved residues E132^3.49^ and R133^3.50^ of the D[E]RY motif form an ionic lock that stabilizes the inactive state (Fig. 1d). Upon receptor activation, this ionic lock breaks apart releasing R133^3.50^ for interaction with a G protein^38,39^. Interestingly, S1P_5_ possesses S134^3.51^ in this motif, which is seen in only 6 class A receptors out of 714, compared to a more common residue Y that is present in 66% of class A receptors.

### Ligand-binding pocket

#### Overall structure of the ligand-binding pocket

The co-crystallized ligand ONO-5430608 (4-{6-[2-(1-naphthyl)ethoxy]-1,2,4,5-tetrahydro-3H-3-benzazepin-3-yl} butanoic acid) has been developed within a series of S1P_5_-selective modulators^25^ and characterized as a potent inverse agonist at S1P_5_ in G_i_-protein-mediated cAMP accumulation assay (EC50 = 1.7 nM) (Supplementary Fig. 6 and Table 2). The ligand was modeled in a strong electron density observed inside the ligand-binding pockets of both receptor molecules in the obtained crystal structure (Fig. 2a and Supplementary Fig. 4). The overall architecture of the pocket, shared by other members of the S1PR family, reflects both zwitterionic and amphipathic properties of the endogenous S1P ligand^20,21^. The pocket is occluded on the extracellular side by the N-terminal alpha-helix packed along ECL1 and ECL2, with a small opening between TM1 and TM7 (Fig. 1f), which has been proposed to serve as the entrance gate for lipid-like ligands^20^. The orthosteric ligand-binding pocket in S1P_5_ consists of a polar charged part, composed of residues from the N-terminal helix and extracellular tips of TM2 and TM3 that interact with the zwitterionic headgroup of S1P, as well as a hydrophobic cavity, lined up by hydrophobic and aromatic residues, which accommodates the alkyl tail of S1P (Fig. 2b). The negatively charged butanoic acid group of ONO-5430608 occupies the polar part of the pocket mimicking the phosphate group of S1P, the core tetrahydro-benzazepine rings fill in space in the middle of the pocket, while the naphthyl-ethoxy group unexpectedly swings over and extends into a previously unidentified allosteric sub-pocket. The sub-pocket is surrounded by non-conserved residues from TM1, TM2, and TM7 and opened in our structure due to a rotameric switch of Y89^2.57^ compared to structures of other S1P receptors (Fig. 2a,b). The distinct amino acid composition of this allosteric sub-pocket suggests that it can serve as a selectivity determinant for S1P_5_-specific ligands and makes the hallmark of the structure described in this work.

**Fig. 2.**
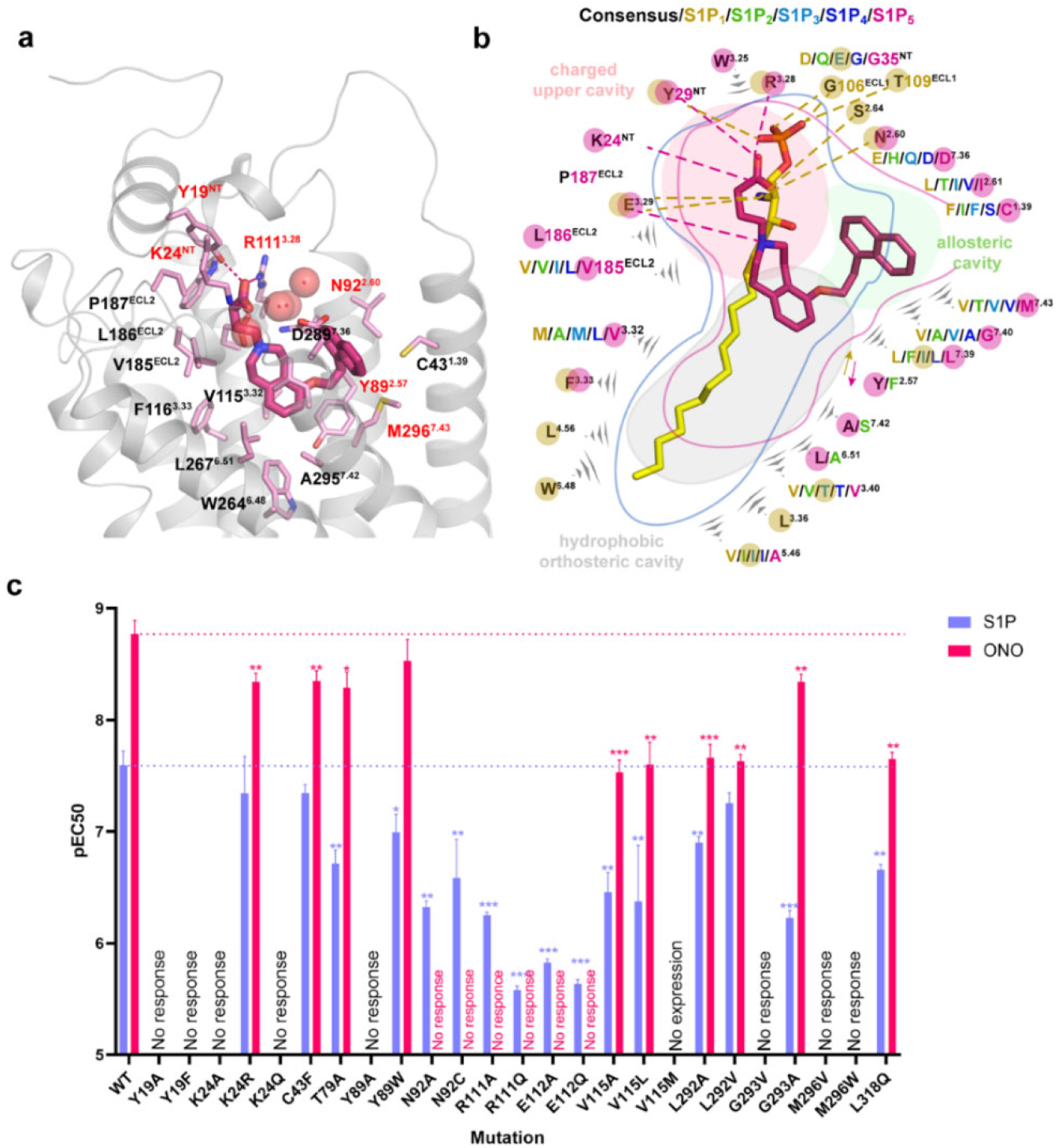
Structural and functional comparison between ONO-5430608 and S1P binding. **a** Binding pose of ONO-5430608 (pink, thick sticks) in S1P_5_ and its interactions with the receptor residues (light pink, thin sticks). Polar interactions are shown as dashed lines. Residues which had mutations disrupting response to ONO-5340608 are labeled in red. **b** Schematic diagram of the ligand binding pocket and interactions between ONO-5430608 and S1P_5_ (this work, PDB ID 7YXA) compared to interactions between S1P and S1P_3_ (PDB ID 7C4S). Residues are color coded according to different S1P receptor subtypes (S1P_1_ - yellow, S1P_2_ - light green, S1P_3_ - light blue, S1P_4_ - dark blue, S1P_5_ - pink). Black stands for the consensus residue shared by all receptors. Residues interacting with ONO-5430608 are highlighted with pink circles, residues interacting with S1P are highlighted with yellow circles. Polar interactions are shown as dashed lines. **c** Potencies (pEC50) of S1P (purple, agonism) and ONO-5430608 (pink, antagonism) at WT and mutants of S1P_5_ in G_i_ protein-mediated signaling assays. Data are shown as mean ± s.d. of 3 independent experiments conducted in triplicates. Data were analyzed by two-sample t-test; * 10^-2^ ≤ *p* < 5 · 10^-2^, ** 10^-3^ ≤ *p* < 10^-2^, *** *p* < 10^-3^. Dose-response curves are shown in Supplementary Fig. 6.

### Functional characterization of the ligand binding hotspots in S1P_5_

To validate the observed ligand binding pose and further expand our knowledge about the ligand selectivity and relative importance of specific residues, we tested 25 structure-inspired ligand-binding pocket mutants of S1P_5_ by a BRET-based cAMP production assay using the endogenous agonist S1P and the co-crystallized inverse agonist ONO-5430608 (Fig. 2c, Supplementary Table 2, and Supplementary Fig. 7). In line with the binding pocket structure description given above, we consequently characterize important interactions in each part.

#### Polar part of the orthosteric pocket

The polar upper part of the binding pocket is highly conserved among the whole S1PR family (Fig. 2b). It consists of residues Y19^N-Term^, K24^N-Term^, N92^2.60^, R111^3.28^, and E112^3.29^ and accommodates the phosphate and primary amine groups of S1P. The receptor’s potential for multiple polar interactions in this region is utilized in anchoring zwitterionic groups of synthetic ligands of S1P receptors. Thus, in our S1P_5_ structure, the carboxyl group of ONO-5430608 is stabilized by polar interactions with Y19^N-term^, K24^N-term^, and R111^3.28^, while the protonated tertiary amine group makes a salt bridge with E112^3.29^, similar to interactions of the phosphate and secondary amine groups of ML056 in S1P_1_ structure^20^. The zwitterionic headgroup of the endogenous S1P ligand bound to S1P_3_ is shifted towards TM1, while retaining the same interactions except for the N-terminal K27^21^.

The mutations disrupting polar interactions with zwitterionic ligand head groups: Y19^N-Term^A/F, K24^N-Term^ A/Q, N92^2.60^A/C, R111^3.28^A/Q, and E112^3.29^A/Q either fully abolish or significantly (by an order of magnitude or more) decrease the response for both ONO-5340608 and S1P (Fig. 2c and Supplementary Table 2). Notably, some mutations have different effects on S1P and ONO-5430608. While mutations of N92^2.60^, R111^3.28^, and E112^3.29^ completely eliminate response to the inverse agonist, they only decrease the potency for S1P. A similar effect of mutations of homologous amino acids on S1P potency was previously observed for S1P_3_^21^. In this case, each of the three amino acids independently interacts with the amine group of S1P (see PDB ID 7C4S). On the other hand, in our S1P_5_-ONO-5430608 structure, these three amino acids are interconnected and form a stable cluster which further interacts with the tertiary amine and the carboxyl group of ONO-5430608. Thus, mutations of any of the three amino acids in S1P_5_ would only partially perturb S1P complex, while they would disrupt the cluster and completely eliminate the binding of ONO-5430608.

Although the locations of residues, known to interact with the phosphate group of S1P from either functional or structural data, are largely conserved between S1P receptors, the effects of their mutations on S1P potency are different. Namely, mutations of N-terminal Y29/19 and K34/24 to alanine render S1P_1_ / S1P_5_, respectively, non-responsive to S1P^20^, while corresponding mutations preserve the interaction with S1P_3_^21^. These data suggest a different orientation of the phosphate headgroup of S1P within the binding pocket in different receptors.

#### Hydrophobic part of the orthosteric pocket

The hydrophobic part of the orthosteric binding pocket in S1P receptors accommodates the lipidic tail of the endogenous ligand or its synthetic analogs such as ML056^20,21,40^. The residues on its bottom are conserved among S1PRs (Fig. 2b) and well characterized^21,40^. The top part of the hydrophobic subpocket in S1P_5_, which in our structure accommodates the tetrahydro-benzazepine double ring system of ONO-5430608, consists of residues V115^3.32^, L292^7.39^, and Y89^2.57^ that are less characterized, although they play an important role in ligand binding.

In our functional assays, mutations of V115^3.32^ to A and L decrease potencies of both S1P and ONO-5430608 (Fig. 2c and Supplementary Table 2). Additionally, Y89^2.57^A abolishes the functional response of both ligands, while Y89^2.57^W preserves it, suggesting that an aromatic residue is crucial at this position.

#### Allosteric sub-pocket

Although ONO-543060 shares a similar zwitterionic headgroup with other co-crystallized S1PR ligands, its hydrophobic tail is substantially different. The bulky naphthyl group of ONO-543060 does not fit well in the relatively narrow hydrophobic cleft of the orthosteric pocket and instead accommodates a previously uncharacterized allosteric subpocket between TM1 and TM7 (Fig. 2a,b). The subpocket is formed by non-conserved hydrophobic residues C43^1.39^ (90 Å^2^ occluded area), I93^2.61^ (127 Å^2^), L292^7.39^ (126 Å^2^), G293^7.40^ (50 Å^2^), and M296^7.43^ (123 Å^2^). Site-directed mutagenesis of residues in the allosteric pocket and functional data suggest a strong role of TM7 residues of the pocket in ligand binding. In particular, mutations L292^7.39^A/V and M296^7.43^V/W decrease ONO-543060 potency by over an order of magnitude (Fig. 2c). On the other hand, mutations C43^1.39^F and G293^7.40^A show almost no effect on either ligand binding. The strengths of the effects appear to correlate with the occluded areas of residues interacting with ONO-543060, as calculated from the crystal structure.

#### Meta molecular dynamics simulations of Y^2.57^ conformational flexibility

The allosteric subpocket displays a large variability in its residues among S1P receptors (Fig. 2b), likely contributing to the exceptional selectivity of ligands targeting it. Interestingly, this pocket is present in our S1P_5_ structure largely due to the flip of one amino acid, Y89^2.57^, compared to other S1PR structures. We thus explored the conformational flexibility of Y^2.57^ in the available structures of S1P_1_, S1P_3_, and S1P_5_ receptors using an enhanced molecular dynamics simulation technique, originally developed by Laio and Parrinello^41^ and known as metadynamics (metaMD), as well as by targeted mutagenesis.

MetaMD facilitates sampling of the free energy landscape along the selected reaction coordinate(s), e.g., a torsion angle, by adding biasing potentials (most commonly positive Gaussians) driving the system out of local minima. By adding multiple Gaussians, the system is discouraged to return to already sampled regions of the configurational space what eventually allows it to escape free energy minima. The free energy landscape can be then recovered as the opposite of the cumulative biasing potential. Here, we used metaMD to estimate free energy profiles along the reaction coordinate corresponding to the torsion rotation of the Y^2.57^ side chain.

The free energy profile of the Y89^2.57^ side chain torsion in S1P_5_ features two minima (Fig. 3a,c): the global minimum corresponds to a downward orientation of Y89^2.57^ as observed in our crystal structure, while the second minimum at a higher energy level is close to an upward orientation of Y^2.57^ found in the S1P_1_ and S1P_3_ crystal structures. On the other hand, the free energy profile of the Y^2.57^ side chain torsion in both S1P_1_ and S1P_3_ has only a single minimum near their crystallographic upward conformations (Fig. 3c). The downward orientation of Y^2.57^ in the latter case is likely hampered by steric clashes with M^3.32^/V^7.43^, making this conformation energetically unfavorable. S1P_5_ has a smaller valine in position 3.32, which does not interfere with the downward orientation of Y^2.57^. At the same time, a more flexible methionine in position 7.43 may swap positions with Y89^2.57^ allowing the latter to switch between the upward and downward conformations.

**Fig. 3.**
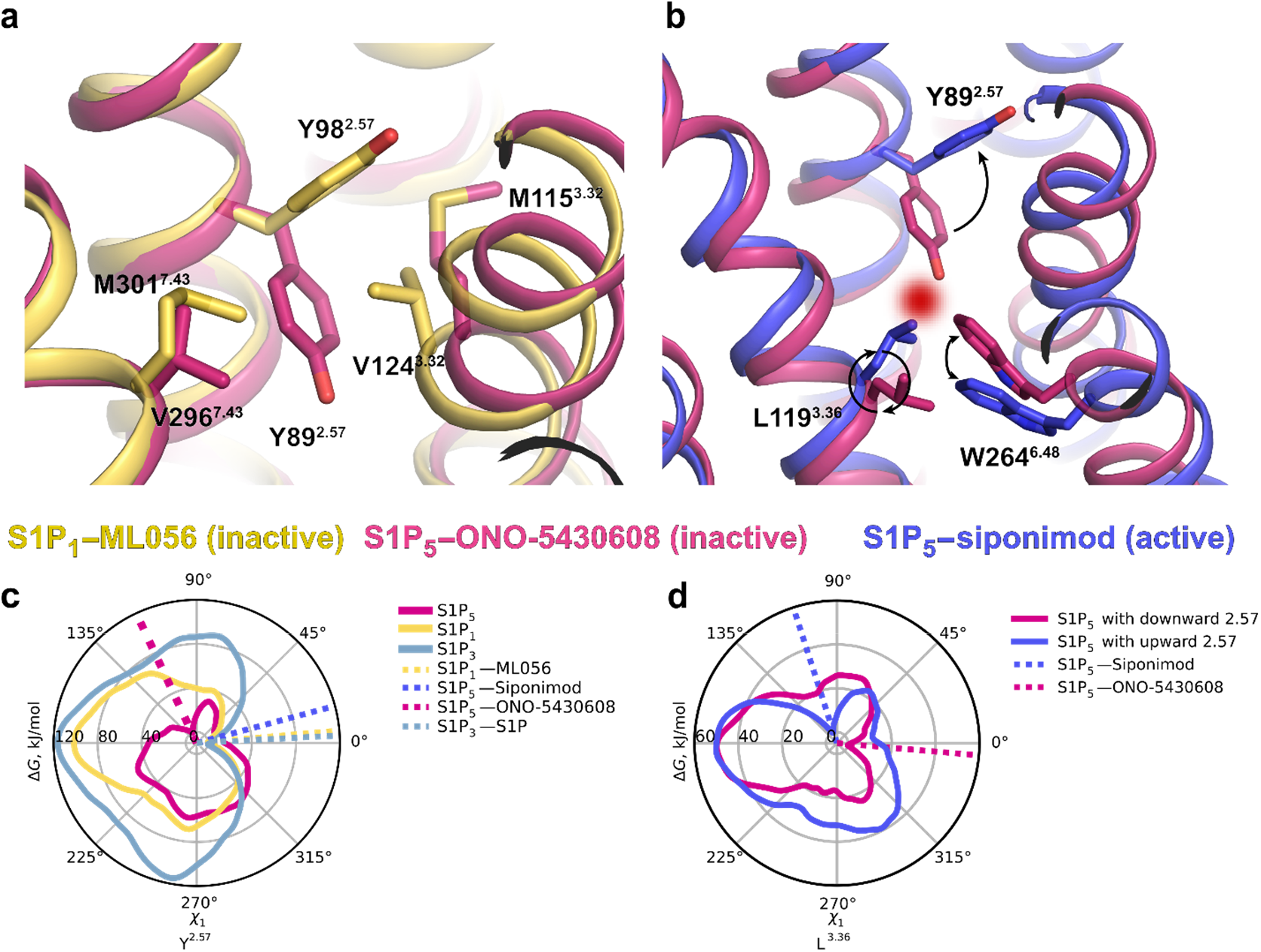
Conformational flexibility of Y^2.57^ and its effect on inverse agonism in S1P receptors. **a** Two distinct upward and downward conformations of Y^2.57^ as observed in crystal structures of S1P_1_-ML056 (PDB ID 3V2Y) and S1P_5_-ONO-5430608 (this work, PDB ID 7YXA), respectively. **b** The downward orientation of Y^2.57^ is incompatible with the active state of the dual toggle switch L^3.36^ – W^6.48^ because of a steric clash. S1P_5_-ONO-5430608 (this work, PDB ID 7YXA, inactive state) is shown in pink and S1P_5_-siponimod (PDB ID 7EW1, active state) is shown in purple. **c** Free energy profiles of the Y^2.57^ side chain torsion angle *χ*_1_ as calculated by metaMD for S1P_1_, S1P_3_ and S1P_5_. Dotted lines correspond to Y^2.57^ conformations in corresponding experimental structures. **d** Free energy profiles of the L^3.36^ side chain torsion angle in S1P_5_ with two alternative orientations of Y^2.57^ as calculated by metaMD. Dotted lines correspond to L^3.36^ conformations in corresponding experimental structures.

#### Insights from molecular docking

To further explore structure-activity relationships from both ligand and protein points of view, we performed molecular docking calculations. To assess the importance of the Y89^2.57^ conformation in ligand binding, we compared two S1P_5_ models: the crystal structure (Y89^2.57^ in the downward conformation) and a metaMD snapshot (Y89^2.57^ in the upward conformation) in their ability to predict binding of ONO-5430608 ligand series^25^. For the crystal structure, the docking scores correlate well with the ligand affinity: the most potent group ‘A’ ligands (IC_50_ between 1 and 100 nM) have docking scores of −37±5, whereas the least potent group ‘C’ ligands (IC_50_ between 1 and 3 μm) have scores −23±4, and for the intermediate group ‘B’ (IC_50_ between 100 nM and 1 μM) scores are −32±5 (Supplementary Fig. 7b). The docking poses of group ‘A’ compounds closely resemble the ligand pose in the crystal structure (Supplementary Fig. 7a). Namely, the interactions of the negatively-charged headgroup with Y19/K24, as well as the interaction of the positively-charged amino group with E112^3.29^, and the position of the double-ring system are preserved. For the upward conformation of Y89^2.57^, the docking scores show no correlation with the ligand affinity, and the docking poses of the group “A” ligands show no consistency between each other and the obtained data from the mutation screening (Supplementary Fig. 7c,d).

Notably, the SAR data for the ONO-5430608 ligand series suggest a role of the substituent position on the core double-ring system in the ligand binding. Namely, most of the lower affinity ligands (group “C”) have a tetrahydroisoquinoline or tetrahydronaphthalene scaffold instead of the tetrahydro-benzazepine, which is more common among group “A” and “B” ligands. Likely, the affinity drop occurs due to the overall ligand shape, rather than the ring size. Namely, most ligands with both substituents placed on the same side of the middle plane across the double-ring system display higher affinity, while all ligands with two substituents placed on different sides have a low affinity (Supplementary Fig. 8), with the only exception, Example 18-2, which however has an amine group placed within the isoquinoline system, compared to other same-side substituents ligands. This notion also suggests a common framework for designing S1P_5_-selective ligands.

#### Inverse agonism

It has been shown that S1P_5_ exhibits a relatively high level of basal activity^42^, while our functional assay revealed that ONO-5430608 acts as an inverse agonist for the G_i_-protein-mediated signaling pathway, reliably decreasing the basal activity level detected by the BRET-based cAMP sensor (Supplementary Fig. 6).

Our S1P_5_ structure in complex with the inverse agonist ONO-5430608 along with previously reported agonist and antagonist-bound structures of S1P_1_,3,5 shed light on the mechanism of inverse agonism. Specifically, the above-mentioned conformational flexibility of Y89^2.57^ may provide a structural background for the basal activity of S1P_5_. As shown by metaMD, the upward orientation of Y89^2.57^ is compatible with both active and inactive conformations of the dual toggle switch L^3.36^-W^6.48^, while the downward orientation of Y89^2.57^ selects the inactive conformation (Fig. 3d). The dual toggle switch is found in the previously reported active state structures of S1P_5_-siponimod as well as in S1P_1_^23^ and S1P_3_^21^ agonist-bound complexes. It induces activation of the P-I-F-motif and an outward movement of the intracellular part of TM6 resulting in G-protein signaling cascade. On the other hand, the dual toggle switch is observed in the inactive conformation in our S1P_5_-ONO-5430608 structure and in the previously published antagonist-bound S1P_1_^20^. The inverse agonist ONO-5430608 induces the downward conformation of Y89^2.57^ that opens the allosteric subpocket and suppresses the switching of L119^3.36^ locking the dual toggle switch in the inactive state (Fig. 3b). Therefore, conformational flexibility of Y89^2.57^ in S1P_5_ provides structural basis for both receptor subtype selectivity and inverse agonism.

### Naturally occurring mutations in S1P_5_

In order to characterize additional functionally important residues in S1P_5_, we performed mapping of known point mutations from genomic databases onto the crystal structure (Supplementary Fig. 9). Multiple databases carry information about S1P_5_ point mutations including gnomAD (229 SNVs)^43^, which contains genomic information from unrelated individuals, and COSMIC (124 point mutations)^44^, which accumulates somatic mutations in cancer. The most frequent gnomAD mutation L318^8.55^Q in helix 8 (3% of the population) was shown to impair G12 signaling^45^; however, according to our functional data it only slightly decreases the potency of S1P in G_i_ mediated signaling (Fig. 2c). It was previously proposed^45^ that a possible cause of this mutation on the signaling impairment is the prevention of palmitoylation of the downstream C322^8.59^ or C323^C-term^. A concomitant cause might be a shift in the helix 8 position due to the loss of a hydrophobic contact between the mutated residue L318^8.55^ and the membrane.

Several individuals have missense mutations in the ligand binding pocket; for example, 3 out of 235,080 samples^43^ contain R111^3.28^L mutation possibly affecting the contacts with the zwitterionic ligand headgroup (Fig. 2). Mutation of another headgroup recognition residue, E112^3.29^G, is less frequent (1 of 234,568). As shown in our functional data, mutations of both of these residues to neutral ones disrupt response to ligands (Fig. 2c). Additionally, two mutations located side-by-side in the putative ligand entrance gateway, C43^1.39^F and M296^7.43^V, are present in the population^43^ at 10^-6^ frequencies. While C43^1.39^F shows little effect in our functional tests, M296^7.43^V disrupts G_i_ signaling response for both S1P and ONO-5430608 (Fig. 2c). Another conserved in S1P receptors, except for S1P_2_, residue A295^7.42^ has a hydrophobic contact with the ligand (Fig. 2a), which becomes altered in case of the A295^7.42^S mutation. Mutation of A295^7.42^S may also directly influence the state of the toggle switch (L119^3.36^-W264^6.48^ in S1P_5_) and may interfere with protein activation, as it was shown for several other receptors, e.g., β_2_-adrenergic receptor^46^ and CCR5^47^. One of the key residues in the sodium binding site, N298^7.45^, has several variations in population: S, D, or K. While the effects of N298^7.45^D and N298^7.45^S are unclear, N298^7.45^K would mimic sodium binding, stabilizing the inactive state of the receptor^48^.

Somatic mutations appearing in COSMIC and not found in the population may be linked to severe cancer impairments. For example, S125^3.42^R disrupts the conservative hydrogen-bond network involving S77^2.45^ and W159^4.50^, destabilizing contacts between TMs 2, 3, and 4^49^ and, likely, disturbing the 7TM fold due to the introduction of a charged residue in a mostly hydrophobic environment.

### Comparison with AlphaFold predictions

Recently, a redesigned artificial intelligence-based protein structure-predicting system AlphaFold v.2^50^ achieved an outstanding breakthrough in approaching the accuracy in protein structure modeling, previously available only from experimental methods. AlphaFold-based approaches started to find multiple applications in structural biology^51^, however, their full capacity and limitations remain to be uncovered. Here, we evaluated the ability of AlphaFold to predict structural features responsible for receptor selectivity and inverse agonism in the S1PR family. For that, we generated 50 *de novo* AlphaFold models for each of the five S1PRs. Overall, the models demonstrated reasonable correspondence to the available experimental structures; for example, Cα RMSDs in the 7TM region between the S1P_5_ models and the inactive state crystal structure (S1P_5_-ONO-5430608) is 1.3±0.2 Å and the active state structure (PDB ID 7EW1, S1P_5_-siponimod) is 3.0±0.2 Å.

The conformational heterogeneity of Y(F)^2.57^ observed in experimental S1PR structures and metaMD simulations was also well captured by AlphaFold predictions (Supplementary Fig. 10a-f). In all S1P_1_, S1P_3_, and S1P_4_ models, Y^2.57^ has an upward conformation, except for a single S1P_1_ model, in which this residue adopts a downward orientation similar to that previously observed in all-atom MD simulations^52^. Furthermore, 19 out of 50 S1P_5_ models display a downward Y^2.57^ orientation, while all the others have an upward Y^2.57^ orientation. Notably, S1P_2_ is the only receptor, in which Y^2.57^ is replaced with F^2.57^ that adopts a downward conformation in all generated models. The downward orientation of F^2.57^ in S1P_2_, similar to that of Y^2.57^ in S1P_5_, opens the allosteric subpocket, which may be targeted to achieve ligand selectivity.

In all available experimental S1PR structures, the conserved dual toggle switch L^3.36^-W^6.48^ displays either active or inactive conformation. AlphaFold predicted both of these conformations for all receptors except for S1P_4_, in which only the active conformation was present in all models (Supplementary Fig. 10a-e). However, AlphaFold models did not fully reflect the mutual relationship between conformations of Y89^2.57^ and L119^3.36^, as observed by metaMD in S1P_5_. Thus, all AlphaFold-predicted S1P_5_ models cluster into three groups (Supplementary Fig. 10e), including the energetically unfavorable conformation with Y89^2.57^-L119^3.36^ in downward-upward orientations while missing the energetically favorable conformation with Y89^2.57^-L119^3.36^ in upward-downward positions. Consequently, we conclude that the current version of AlphaFold could not consistently generate an S1PR structure in a specific signaling state, sometimes mixing the features of different conformations in a single model. These findings are corroborated by a recent study of several other GPCRs^53^.

One of the most intriguing AlphaFold-related questions is how useful are the predicted models for structure-based drug design^54^. To test it in application to S1PR targets, we constructed three benchmarks, mimicking virtual ligand screening campaigns, and compared the available experimental structures and AlphaFold models by their ability to distinguish high-affinity ligands from low-affinity binders and decoys. Our results demonstrated that crystal structures outperform AlphaFold-generated models in several scenarios (Supplementary Fig. 11). Namely, our S1P_5_ crystal structure showed substantially better overall ranking and top-10% enrichment among both ONO-5430608-like inverse agonists (“ONO” benchmark) and S1P_5_-selective ligands (“Selective” benchmark). In the case of the non-selective ligand benchmark (mostly S1P_1_ agonists), the best performance was achieved for several experimental S1PR structures determined in complex with non-selective ligands, e.g., S1P_1_-siponimod complex (Supplementary Fig. 11), while our S1P_5_ structure fared on par with AlphaFold models.

## DISCUSSION

Here, we present the 2.2 Å crystal structure of the human S1P_5_ receptor in complex with its selective inverse agonist. The structure was obtained by room temperature SFX data collection at PAL-XFEL using sub-10 μm crystals. In combination with site-directed mutagenesis, functional assays, metaMD simulations, and docking studies, this structure revealed molecular determinants of ligand binding and selectivity as well as shed light on the mechanism of inverse agonism in the S1PR family. The obtained structure also allowed us to map locations of known missense SNVs from gnomAD and COSMIC genome databases and annotate their potential functional roles providing future insights in personalized medicine approaches.

We found that the inverse agonist ONO-5430608 binds to the receptor’s orthosteric site, suppressing S1P_5_ basal activity. Highly conserved residues Y19^N-term^, K24^N-term^, R111^3.28^, and E112^3.29^ play an essential role in the recognition of both ONO-5430608 and its native ligand S1P. The naphthyl group of ONO-5340608 occupies an allosteric subpocket that was not previously observed in any other S1PR structure. While the orthosteric site is highly conserved in the S1PR family, the allosteric subpocket is composed of unique residues and is present in our S1P_5_ structure due to the conformational switch of a single residue Y^2.57^. Functionally important residues were revealed by structure-guided site-directed mutagenesis and G_i_ signaling assays. We further used metaMD simulations to explore the conformational flexibility of Y^2.57^ in S1PRs and established its role in receptor subtype selectivity and inverse agonism. The role of Y^2.57^ in binding of selective ligands was also confirmed by comparative molecular docking simulations. Furthermore, taking advantage of the availability of several experimental structures of S1PRs in different functional states, we tested the ability of AlphaFold to predict *de novo* specific conformational states for S1PRs and to provide reliable templates for structure-based virtual ligand screening. While the AlphaFold-generated models showed a close similarity to experimental structures and captured conformational diversity of conserved structural motifs, the models did not provide a full description of specific signaling states and showed subpar performance in virtual ligand screening compared to experimental structures.

Our structure along with our functional and computer modeling data may facilitate the rational design of ligands that could further serve as lead or tool compounds for detailed elucidation of biological function of S1P_5_ and therapeutic developments. S1P_5_ is emerging as a promising drug target. Inhibiting S1P_5_ by an inverse agonist could create new therapeutic strategies against neuroinflammation and degeneration where the high ligand selectivity would diminish the off-target effects. While S1P_1_ has a broad expression profile, S1P_5_ is expressed predominantly in brain tissues^8^; thus, a highly selective compound would afford more localized control over associated CNS disorders not affecting peripheral processes in the body.

## METHODS

### Protein engineering for structural studies

The human wild type gene *S1PR5* (UniProt ID Q9H228) was codon-optimized by Genescript for insect cell expression and modified by adding a hemagglutinin signal peptide (HA; KTIIALSYIFCLVFA), a FLAG-tag for expression detection, and an Ala-Gly-Arg-Ala linker at the N-terminus. An apocytochrome b562RIL (BRIL)^26^ was inserted in the third intracellular loop between A223 and R241 to stabilize the receptor and facilitate crystallization. The C-terminus was truncated after Val321, and a PreScission cleavage site was added after it to enable removal of the following 10×His tag used for IMAC purification (Supplementary Fig. 1). The resulting construct was cloned into a pFastBac1 (Invitrogen) plasmid. The full DNA sequence of the S1P_5_ crystallization construct is provided in Supplementary Table 3.

### Protein expression

Using the Bac-to-Bac system (Invitrogen), a high titer (10^9^ particles per ml) virus encoding the crystallization construct was obtained. Sf9 (Novagen, cat. 71104) cells were infected at a density (2-3)×10^6^ cells per ml and a multiplicity of infection (MOI) 4-8, incubated at 28 °C, 120 rpm for 50-52 h, harvested by centrifugation at 2,000 g and stored at −80 °C until further use.

### Protein purification

Cells were thawed and lysed by repetitive washes (Dounce homogenization on ice, and centrifugation at 128,600 g for 30 min at 4°C) in hypotonic buffer (10 mM HEPES pH 7.5, 20 mM KCl, and 10 mM MgCl_2_) and high osmotic buffer (10 mM HEPES pH 7.5, 20 mM KCl, 10 mM MgCl_2_, and 1 M NaCl) with an addition of protease inhibitor cocktail [PIC; 500 μM 4-(2-aminoethyl)benzenesulfonyl fluoride hydrochloride (Gold Biotechnology), 1 μM E-64 (Cayman Chemical), 1 μM leupeptin (Cayman Chemical), 150 nM aprotinin (A.G. Scientific)] with the ratio of 50 μl per 100 ml of lysis buffer. Membranes were then resuspended in 10 mM HEPES pH 7.5, 20 mM KCl, 10 mM MgCl_2_, 2 mg/ml iodoacetamide, PIC (100 μl per 50 ml of resuspension buffer), and 50 μM ONO-5430608 (4-{6-[2-(1-Naphthyl)ethoxy]-1,2,4,5-tetrahydro-3H-3-benzazepin-3-yl}butanoic acid; Example 18(18)^25^, received as a gift from Ono Pharmaceutical) for 30 min at 4 °C and then solubilized by addition of 2× buffer (50 mM HEPES, 500 mM NaCl, 2 %w/v n-dodecyl-β-D-maltopyranoside (DDM; Anatrace), 0.4 %w/v cholesteryl hemisuccinate (CHS; Sigma), 10 %v/v glycerol) and incubation for 3 h at 4 °C with 10 rpm rotation. All further purification steps were performed at 4 °C. The supernatant was clarified by centrifugation (292,055 g, 60 min, 4°C) and bound to 2 ml of TALON IMAC resin (Clontech) overnight with 10 rpm rotation in the presence of 20 mM imidazole and NaCl added up to 800 mM. The resin was then washed with ten column volumes (CV) of wash buffer I (8 mM ATP, 50 mM HEPES pH 7.5, 10 mM MgCl_2_, 250 mM NaCl, 15 mM imidazole, 50 μM ONO-5430608, 10 %v/v glycerol, 0.1/0.02 %w/v DDM/CHS), then with five CV of wash buffer II (50 mM HEPES pH 7.5, 250 mM NaCl, 50 mM imidazole, 50 μM ONO-5430608, 10 %v/v glycerol, 0.5/0.01 %w/v DDM/CHS), then eluted with (25 mM HEPES pH 7.5, 250 mM NaCl, 400 mM imidazole, 50 μM ONO-5430608, 10 %v/v glycerol, 0.05/0.01 %w/v DDM/CHS) in several fractions. Fractions containing target protein were desalted from imidazole using PD10 desalting column (GE Healthcare) and incubated with 50 μM ONO-5430608 and a His-tagged PreScission protease (homemade) overnight with 10 rpm rotation to remove the C-terminal 10×His tag. Protein was concentrated up to 40–60 mg/ml using a 100 kDa molecular weight cut-off concentrator (Millipore). The protein purity was checked by SDS-PAGE. Yield and monodispersity were estimated by analytical size exclusion chromatography. Stability and stabilizing effect of the ligand were measured by microscale fluorescent thermal stability assay^55^ (Supplementary Fig. 2).

### Thermal Stability Assay

Microscale fluorescent thermal stability assay^55^ was conducted using a CPM dye (7-Diethylamino-3-(4-maleimidophenyl)-4-methylcoumarin, Invitrogen) dissolved in DMF at 10 mM. This CPM stock solution was diluted to 1 mM in DMSO and then added to working buffer at 10 μM. 1 μg of the target protein was added to 50 μL of working buffer (25 mM HEPES, 250 mM NaCl, 10 %v/v glycerol, 0.05 %w/v DDM, 0.01 %w/v CHS) with CPM, and the melting curve was recorded on a Rotor-Gene Q real-time PCR cycler (Qiagen) using a temperature ramp from 28 to 98°C with 2°C/min rate. The fluorescence signal was measured in the Blue channel (excitation 365 nm, emission 460 nm), and the melting temperature was calculated as the maximum of the fluorescence signal derivative with respect to temperature.

### LCP crystallization

Purified and concentrated S1P_5_ was reconstituted in LCP, made of monoolein (Nu-Chek Prep) supplemented with 10 %w/w cholesterol (Affymetrix), in 2:3 (v/v) protein:lipid ratio using a syringe lipid mixer^27^. The obtained transparent LCP mixture was dispensed onto 96-wells glass sandwich plates (Marienfeld) in 40 nl drops and covered with 900 nl precipitant using an NT8-LCP robot (Formulatrix) to grow crystals for synchrotron data collection. To prepare crystals for XFEL data collection, the protein-laden LCP mixture was injected into 100 μl Hamilton gas-tight syringes filled with precipitant as previously described^27^. All LCP manipulations were performed at room temperature (RT, 20–23 °C), while plates and syringes were incubated at 22 °C. Crystals of S1P_5_ grew to their full size of < 30 μm (in plates) or < 10 μm (in syringes) within 3 days in precipitant conditions containing 100–300 mM KH_2_PO_4_ monobasic, 28–32 %v/v PEG400, and 100 mM HEPES pH 7.0.

### Diffraction data collection and structure determination

#### Synchrotron data collection

Synchrotron data for S1P_5_-ONO-5430608 crystals were collected at the PX1 beamline of the Swiss Light Source of the Paul Scherrer Institute (Villigen, Switzerland). Data were collected at the 1.0 Å wavelength using an Eiger 16M detector with the beam size matching the crystal size, as seen by the initial 2D mesh scan. The data collection was performed using manual X-ray crystal centering: a 2D mesh scan followed by a 1D mesh scan, selecting the best diffracting spots by visual inspection. The centering was followed by data collection of 90° with 0.5° oscillation and a total doze of 30 MGy.

#### Data collection using X-ray free electron laser

XFEL data for S1P_5_-ONO-5430608 crystals were collected at the NCI (Nanocrystallography and Coherent Imaging) beamline of the Pohang Accelerator Laboratory X-ray Free Electron Laser (PAL-XFEL), Pohang, South Korea. The PAL-XFEL was operated in SASE mode at the wavelength of 1.278 Å (9.7 keV) and 0.2% bandwidth, delivering individual X-ray pulses of 25-fs duration focused into a spot size of 2×3 μm using a pair of Kirkpatrick-Baez mirrors. LCP laden with dense suspension of protein microcrystals was injected at room temperature inside a sample chamber filled with helium (23 °C, 1 atm) into the beam focus region using an LCP injector^56^ with a 50-μm-diameter capillary at a flow rate of 0.15 μl/min. Microcrystals ranged in size from 5 to 10 μm. Diffraction data were collected at a pulse repetition range of 30 Hz with a Rayonix MX225-HS detector, operating in a 4×4 binning mode (1440×1440 pixels, 30 fps readout rate). The beam was not attenuated and delivered full intensity (5×10^11^ photons per pulse). A total number of 490,000 detector images were collected. Due to a high systematic background, Cheetah^57^ was initially used only to apply dark current calibration, and all images were used for further processing. The overall time of data collection from a sample with a total volume of about 36 μl was approximately 4 hours and yielded 6,918 indexed frames with 7,492 crystal lattices.

#### XFEL data processing with CrystFEL

During the XFEL data collection, a high systematic background scattering from upstream to the interaction point occurred due to a high intensity X-ray lasing conditions (Supplementary Fig. 4), which prevented from establishing suitable Cheetah hit finding parameters during the beamtime and complicated the overall data processing. All data processing was performed using CrystFEL 0.8.0^58^. Here we describe steps that we took to improve data quality as much as possible starting from the available data with a high background level. For all CrystFEL runs (Supplementary Table 4), peak search was limited with max-res=340, min-res=50 to search for peaks in the region between the beamstop and the LCP ring, and the frames were limited to a 12,000 subset of all frames, selected with minimum 5 peaks with SNR 2.7. Initially, typical starting peak finding parameters (SNR=5.0, threshold=100) in CrystFEL were used for data processing, yielding only 2,036 crystals with indexing=mosflm,dirax,xgandalf (Supplementary Table 4 column A). Initial peak search parameter adjustment, as described in CrystFEL tutorial^58^, led to the value of SNR=2.7 and threshold=30, which yielded 5,275 crystals (Supplementary Table 4 column B). Applying --median-filter=5 allowed to further increase the number of crystals to 7,189, while increasing SNR to 4.0 (Supplementary Table 4 column C).

Spot integration parameters had the biggest impact on the merged data quality. First, changing spot integration model from rings-nograd model, that assumes flat background around a spot, to rings-grad, that performs 2D-fitting of each spot background profile, decreased overall R_split_ from 29.7 % to 19.4 % (Supplementary Table 4 column D) and increased the highest resolution shell CC* from 0.618 to 0.666 Second, increasing local-bg-radius from 3 to 5, and using int-radius=3,5,8 instead of default 4,5,8 further improved data quality with the highest resolution shell CC* equal to 0.716 (Supplementary Table 4, columns E-F). The final merging was performed with partialator, iterations=2, push-res=5.0, and model=ggpm (Supplementary Table 1).

#### Structure determination and refinement

The structure was initially solved by molecular replacement using phenix.phaser^59^ with two independent search models of the poly-alanine S1P_1_ 7TM domain (PDB ID 3V2Y) and BRIL from the high-resolution A2AAR structure (PDB ID 4EIY). Model building was performed by cycling between manual inspection and building with Coot^60^ v. 0.9.6 using both 2mFo-DFc and mFo-DFc maps and automatic refinement with phenix.refine^61^ v. 1.19.2 using automatic torsion-angle NCS restraints and 2 TLS groups. Ligand restraints were generated using the web server GRADE v. 1.2.19 (http://grade.globalphasing.org). The S1P_5_ structures from two molecules A and B in the asymmetric unit show very high similarity (Ca RMSD 1.0 Å within 7TM; 1.3 Å all-atom RMSD). The main difference includes flexible ECL1 and conformations of several side chains exposed to the lipid bilayer and solvent. The final data collection and refinement statistics are shown in Supplementary Table 1. The relatively high Rf_re_e of the structure can be explained by high systematic background scattering and the presence of a pseudotranslation-related modulation in the diffraction intensities. The modulation is produced by the NCS operator (x, y, z) → (1/8 + x, -y, -z) as seen in the Patterson maps of both single-crystal and XFEL data: ~0.7 and 0.3 origin peak high at (3/8, 1/2, 0) for single-crystal and serial data, respectively.

### AlphaFold predictions

Prediction runs were executed using AlphaFold^50^ v. 2.1.1+110948 with a non-docker setup (https://github.com/kalininalab/alphafold_non_docker, git commit 7ccdb7) and an updated run_alphafold.sh wrapper with added --random-seed parameter. The use of templates was disabled by setting “max_template_date” to 1900-01-01. 50 AF2-models (ranked_….pdb models) were generated for each of 5 human S1P receptors with protein sequences obtained from UniProt. For each receptor, 10 prediction runs with different seeds (--random-seed”=<run_number>) were executed; each run generated 5 models. Structures were used as provided by the Alphafold’s pipeline with Amber relaxation (see Suppl. Methods 1.8.6 in Ref. 51 for details) without any further modifications.

### MD simulations

Molecular dynamics simulations were conducted for the wild type human S1P_1_, S1P_3_, and S1P_5_ receptors based on the X-ray structures 3V2Y^20^ (residues V16-K300), 7C4S^21^ (G14-R311), and the structure reported in the present study (S12-C323), respectively. All engineered mutations were reverted back to the WT amino acids, and all missing fragments were filled using MODELLER^62^. Receptors were embedded into lipid bilayers consisting of 1-palmitoyl-2-oleoyl-sn-glycero-3-phosphatidylcholine (POPC) lipids and solvated with TIP3P waters and Na^+^/Cl^-^ ions (to guarantee the electroneutrality of the systems and the ionic strength of 0.15 M) by means of the CHARMM-GUI web-service^63^. The obtained in this way starting models (with 61,666/61,763/56,303 atoms including 119/117/123 POPC molecules in the S1P_1_/S1P_3_/S1P_5_ systems, respectively) were subject to standard CHARMM-GUI minimization and equilibration protocol, i.e., the steepest descent minimization (5,000 steps) was followed by a series of short equilibration simulations in the NPT ensemble using Berendsen thermostat and barostat with the restraints on protein and lipids gradually released.

In order to estimate free energy profiles along the rotation of the *χ*_1_ torsion angle in the side chain of Y^2.57^, we employed the metadynamics (metaMD) approach^41^, which is based on addition of biasing repulsive potentials (“hills”, typically Gaussians) to the total potential of the system to enhance the sampling of the configurational space along the chosen reaction coordinates. The deposition rate for hills in metaMD simulations was 1 ps^-1^; the width and height of deposited hills were equal to 0.1 rad (~5.7°) and 0.5 kJ/mol, respectively. The metaMD simulations were run for 10 ns each. To test for convergence of the metaMD simulations, we applied the following method^64^: the free energy difference between two regions of the obtained free energy profiles (corresponding to the crystallographic orientations of Y^2.57^) as a function of simulation time were plotted (Supplementary Fig. 12a-c). In case of convergence, this difference should not change with the progress of simulations as the systems diffuse freely along the reaction coordinate.

For the metaMD simulations, Nose–Hoover thermostat and Parrinello–Rahman barostat were used. The temperature and pressure were set to 323.15 K and 1 bar with temperature and pressure coupling time constants of 1.0 ps^-1^ and 0.5 ps^-1^, respectively. All MD simulations were performed with GROMACS^65^ v. 2020.2 using PLUMED plugin^66^ to enable metaMD. The time step of 2 fs was used for all production simulations. The CHARMM36 force field^67^ was used for the proteins, lipids, and ions.

### SAR and molecular docking (with ONO compounds)

For S1P_5_ docking studies, we used chain B from our S1P_5_-ONO-5430608 crystal structure and a metaMD snapshot with the upward conformation of Y89^2.57^. Chain B was selected based on the quality of 2mFo-DFc maps around the ligand and surrounding residues. Molecular docking was performed using ICM Pro v. 3.9-1b (Molsoft, San Diego). We removed ligands and converted the receptor models into an ICM format using default settings, which includes building missing sidechains, adding hydrogens, energy-based Gln/Asn/His conformation optimization, and removal of all water molecules. Same docking box was selected for both models, aligned by their 7TM domains, to encompass both orthosteric and allosteric binding pockets. For each ligand we repeated docking runs 5 times with the effort parameter (ligand sampling depth) set at 16, each time saving 3 best conformations. Ligand structures and their affinities (IC_50_ values from radioligand binding assays) at S1P_5_ receptors were taken from the published patent^25^.

In AlphaFold models analysis, 50 models predicted by the AlphaFold algorithm were compared with both chains of our S1P_5_ crystal structure and other available S1PR crystal structures. All structures were prepared as described above. S1P_5_ ligands from ChEMBL v. 29 were accessed via the web-interface (https://www.ebi.ac.uk/chembl/) using the S1P_5_’s ChEMBL target ID. Ligands were converted to 3D and charged at pH 7.0 using Molsoft ICM. For each model, ligand screening was performed three times with docking effort 1. Three ligand benchmarks (Supplementary Fig. 11) were used: 1. “ONO” series: active molecules from Ref. 25 (group A, 1 nM < IC_50_ < 100 nM), inactive molecules from Ref. 25 (group C, 1 μM < IC_50_ < 3μM) and decoys; 2. “Selective” series: active molecules from Refs. 25,69 (group A or IC_50_ < 100 nM, correspondingly), inactive molecules from Refs. 25,69 (group C or IC_50_ >= 1 μM, correspondingly), and decoys; 3. “Non-selective” series: active molecules from ChEMBL (pChembl > 7.0, mostly S1P_1_ agonists), inactive molecules from ChEMBL (pChembl < 5.0), and decoys. Decoy molecules were selected from the Enamine REAL library [https://enamine.net/compound-collections/real-compounds/real-database], matching the distribution of active molecules by charge and weight. The benchmarks have the following ratios of active:inactive:decoy molecules: 6:5:60 for “ONO”, 12:10:120 for “Selective”, and 158:39:1207 for “Non-selective”, with the imbalance parameter (ratio of the total library size to the number of active molecules in it) of 11.8, 11.8, and 8.8, respectively. For estimation of the virtual screening quality, metrics enrichment at 10% and receiver operating characteristic (ROC) – area under the ROC curve (AUC) were used, as implemented in RDKIT v. 2021-03-4 (http://www.rdkit.org^69^).

### Plasmids for functional assays

The human wild-type *S1PR5* gene (UniProt ID Q9H228) with an N-terminal 3 × HA epitope (YPYDVPDYA) tag was cloned into pcDNA3.1+ (Invitrogen) at KpnI(5’) and XhoI(3’). Point mutations were introduced by overlapping PCR. All DNA sequences were verified by Sanger sequencing (Evrogen JSC). Sequences of all primers used in this work are listed in Supplementary Table 5.

### Cell surface expression determined by ELISA

Cell surface expression of S1P_5_ receptor variants was determined by whole-cell ELISA^70^. Briefly, HEK293T cells were seeded in 24-well cell culture plates (0.2×10^6^ cells in 0.5 ml of medium per well) and transfected separately by 3 μg of each expression plasmids based on pCDNA3.1(+) vector using common Lipofectamine 3000 protocol. After 12–18 h incubation in a CO2 incubator at 37 °C for receptor expression, the cell culture plates were placed on ice, the media was aspirated completely, and the cells were washed once with ice-cold TBS to remove any residual media. Then the cells were fixed using 400 μl of 4 %w/v paraformaldehyde (PFA), followed by three 400–500 μl washes with TBS. After surface blocking with 2 %w/v protease-free BSA (A3059, Sigma) solution in TBS, HRP-conjugated anti-HA antibody high affinity (3F10) (Roche) at a dilution of 1:2000 in TBS + 1 %w/v protease-free BSA and TMB ready-to-use substrate (T0565, Sigma) was used for ELISA procedure. The ELISA results were normalized by Janus Green staining. Cells transfected with empty vectors (pCDNA3.1+) were used to determine background.

### Functional assays with BRET-based cAMP sensor

G_i_-protein-mediated signaling responses to endogenous agonist S1P and inverse agonist ONO-5430608 were assayed for human WT and mutant S1P_5_ receptors using Bioluminescence Resonance Energy Transfer (BRET) based EPAC biosensor^71^. Briefly, transfections were carried out by Lipofectamine 3000 according to standard protocol using HEK293T cells seeded in a 100 mm cell culture plate, receptor cDNA vectors (10 μg each), and EPAC biosensor cDNA vector (10 μg) needed for evaluation of cAMP production. Transfected cells were split into 96-well plates at 10^5^ cells per well and incubated for 16-18 h. To measure response for S1P, 60 μl of PBS was added to each well followed by addition of 10 μl of a 50 μM coelenterazine-h, 10 μl of 300 μM forskolin and 10 μl of 100 μM 3-isobutyl-1-methylxanthine (IBMX) solutions. After 10-min incubation, either 10 μl of vehicle or 10 μl of S1P at different concentrations in 0.5 %w/v fatty acid-free BSA (10775835001, Roche) solution in PBS was added. To measure response for ONO-5430608, 70 μl of PBS was added to each well followed by addition of 10 μl of 50 μM coelenterazine-h and 10 μl of 100 μM IBMX solutions. After 10-min incubation, either 10 μl of vehicle or 10 μl of ONO-5430608 at different concentrations in PBS was added. The plate was then placed into a CLARIOstar reader (BMG LABTECH, Germany) with a BRET filter pair (475±30 nm - coelenterazine-h and 550±40 nm - YFP). The BRET signal was determined by calculating the ratio of the light emitted at 550 nm to the light emitted at 480 nm. The EC50 values were calculated using the three-parameter dose - response curve fit in GraphPad Prism v. 9.3. Three independent experiments were performed in triplicate.

## DATA AVAILABILITY

Data supporting the findings of this manuscript are available from the corresponding authors upon reasonable request. A reporting summary for this article is available as a Supplementary Information file. Coordinates and structure factors have been deposited in the Protein Data Bank (PDB) under the accession code 7YXA. Raw diffraction data have been deposited to CXIDB database (https://www.cxidb.org) under the accession number 196.

## Acknowledgements

We thank S. Ustinova, A. Podzorov, A. Awawdeh, P. Utrobin, and Yu. Semenov for technical assistance, T. Maruyama for providing ONO-5430608. We acknowledge the Paul Scherrer Institut, Villigen, Switzerland for provision of synchrotron radiation beamtime at beamline PX1 of the SLS and would like to thank Dr. V. Olieric for assistance. The authors are grateful to the staff of the accelerator and beamline departments at PAL-XFEL for their technical support. The XFEL experiments were performed at the NCI PAL-XFEL experimental station under the proposal No. 2019-2nd-NCI-012. Protein production and crystallization was supported by the Russian Science Foundation project no. 19-14-00261. Synchrotron data collection and treatment were supported by the Russian Ministry of Science and Higher Education grant No. 075-15-2021-1354. SFX data collection strategy was developed with the support from the Russian Foundation for Basic Research (RFBR, project № 18-02-40020). Functional assays were implemented with the support of the Ministry of Science and Higher Education of the Russian Federation (agreement # 075-00337-20-03, project FSMG-2020-0003). J.P. and Y.C. were supported by the National Research Foundation of Korea (grant No. NRF-2017M3A9F6029736). The authors are grateful to the Global Science Experimental Data Hub Center (GSDC) for data computing and the Korea Research Environment Open NETwork (KREONET) for network service provided by the Korea Institute of Science and Technology Information (KISTI) and to the Data Processing Center of Moscow Institute of Physics and Technology for high-performance data computing infrastructure and technical support. V.C. acknowledges that the University of Southern California is his primary affiliation.

## Author contributions

E.L. and A.Gu. optimized the constructs, developed the expression and purification procedure, expressed and purified the protein, screened the ligands, and crystallized the protein–ligand complexes.

E.M., D.V., A.M. and V.C. collected X-ray diffraction data at PAL XFEL.

J.P. set up the XFEL experiment, beamline, controls, and data acquisition; operated the beamline. H.H., U.W. helped develop and operate the LCP injector. W.L. helped with the XFEL sample preparation. S.P. contributed to the experimental system installation at the beamline. G.P. contributed to DAQ and data handling. H.J.H. contributed to the Rayonix detector installation and operation.

A.Gu., E.M., E.L., K.K and A.M. collected synchrotron data at SLS.

A.Ge. performed and analyzed cell signaling and cell surface expression assays.

E.M., D.V. and V.B. processed diffraction data.

G.B. helped with X-ray diffraction data interpretation and analysis.

E.M., V.B., V.C. performed structure determination and refinement.

V.B., V.C., E.L., A.Gu., E.M., A.M., A.L. performed project data analysis/interpretation.

M.E., A.L., M.S. helped with construct optimization, protein expression and purification.

P.P. advised on the protein construct design.

E.M., M.K. performed molecular docking.

P.O., E.M. performed MD simulations.

E.M., I.G., V.B. performed AlphaFold simulations and their data analysis.

A.L., I.O., P.Kh., A.R., Y.C. helped with experimental work and project organization.

E.L., E.M., A.Gu., P.O., A.M., V.B., V.C., wrote the manuscript with the help from other authors.

V.C, A.M., V.B. and V.G. initiated the project.

A.M. and V.B. organized the project implementation, were responsible for the overall project management, and co-supervised the research.

V.C. supervised the overall project.

## Competing interests

Authors declare no competing interests.

## Notes

### Competing Interest Statement

The authors have declared no competing interest.

