## Supplementary for "Structural basis for receptor selectivity and inverse agonism in S1P_5_ receptors"

### Supplementary Figures

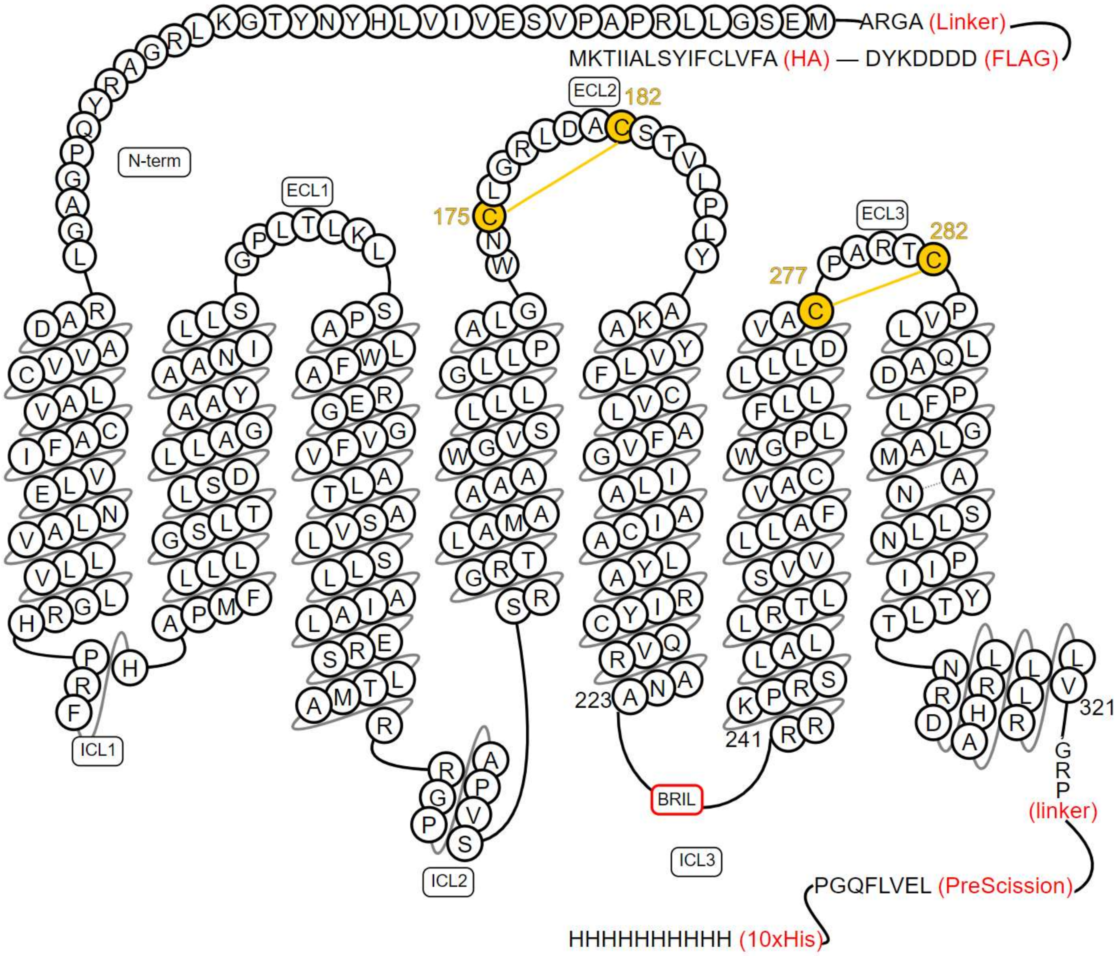


**Supplementary Fig. 1 Crystallization construct of S1P$_5$.** HA signal peptide, FLAG tag, and AGRA linker are attached to the N-terminus; BRIL is inserted in the ICL3 between residues A223 and R241; the C-terminus is truncated at V321 followed by GRP linker, PreScission site, and decahistidine tag.

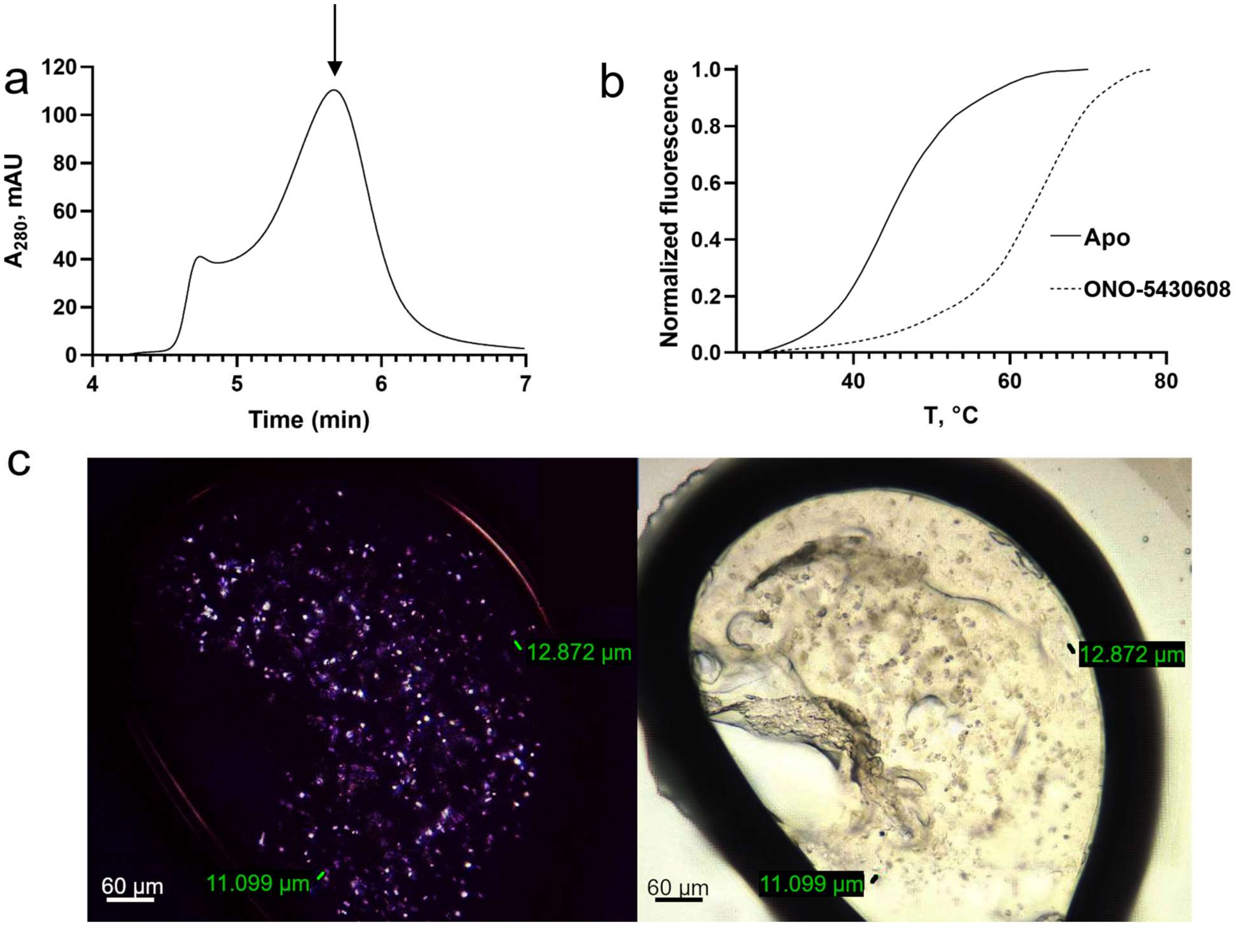


**Supplementary Fig. 2 Characterization and crystallization of S1P$_5$-ONO-5430608. a** Analytical size exclusion chromatography analysis of purified S1P$_5$ in complex with ONO-5430608, showing mostly monomeric protein preparation (the monomer peak is shown with an arrow). **b** Thermal shift assay using CPM fluorescence. ONO-5430608 increases the thermal stability of S1P$_5$ by 19 °C. **c** Crystals of S1P$_5$-ONO-5430608 grown in lipidic cubic phase as visualized by cross-polarized and direct light microscopy.

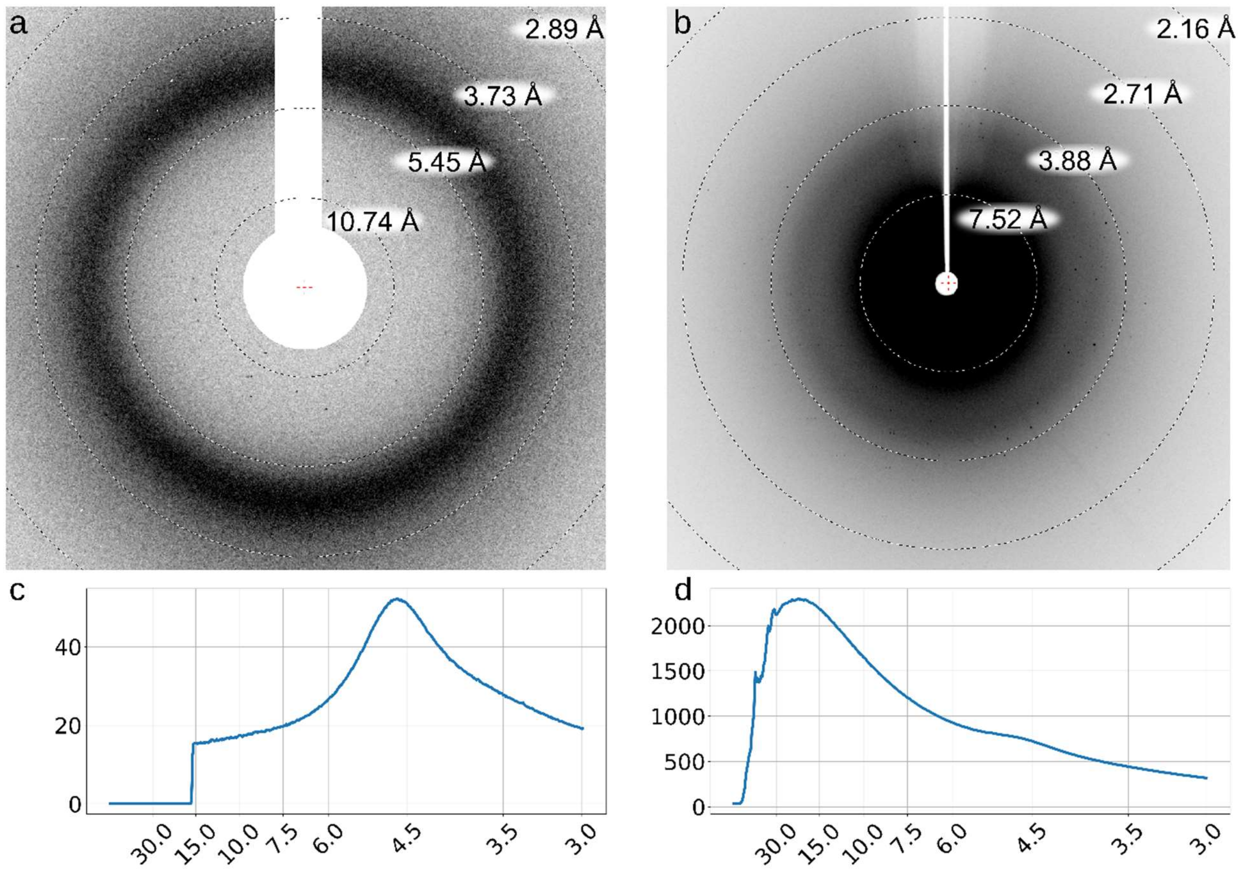


**Supplementary Fig. 3 Examples of diffraction images collected at PAL-XFEL. a,b** Diffraction images. **c,d** radial profiles. Diffraction image (**a**) and its radial profile (**c**) correspond to a typical low background experiment. Diffraction image (**b**) and its radial profile (**d**) correspond to the high background data collection described in this article.

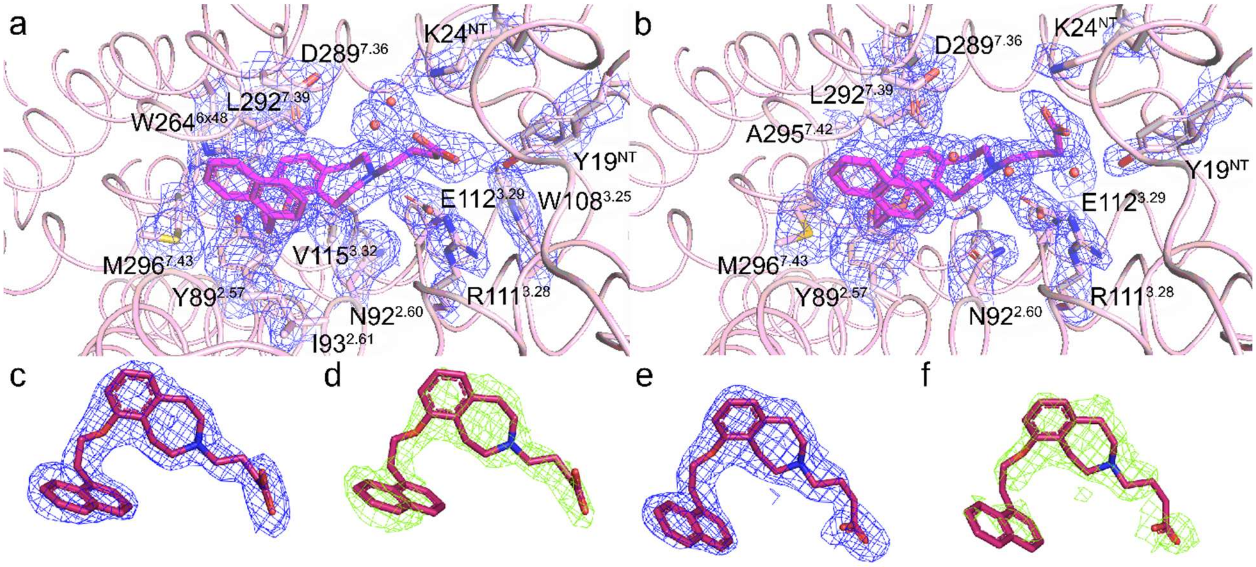


**Supplementary Fig. 4 Examples of electron density. a,b** 2mFo-DFc electron density maps around ONO-5430608 and ligand binding pocket residues and water molecules within 4 Å of the ligand in chains A (**a**) and B (**b**), contoured at 1.0 σ level. **c,e** 2mFo-DFc electron density maps around ONO-5430608 in chains A (**c**) and B (**e**), contoured at 1.5 σ. **d,f** Simulated annealing ligand omit 2mFo-DFc electron density maps around ONO-5430608 in chains A (**d**) and B (**f**), contoured at 1.5 σ.

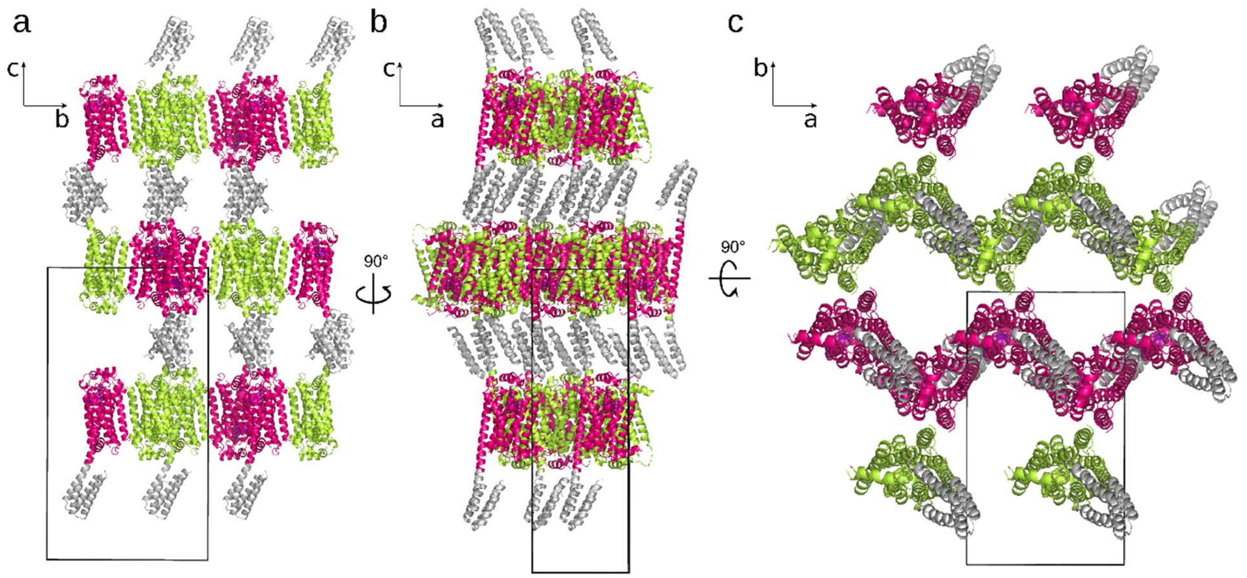


**Supplementary Fig. 5 Crystal packing of S1P$_5$-ONO-5430608. a-c** Crystal packing is shown in three orthogonal orientations. The unit cell is outlined by a black box. S1P$_5$ is shown in pink (chain A) and light green (chain B), BRIL fusion partner in grey, ligand in purple.

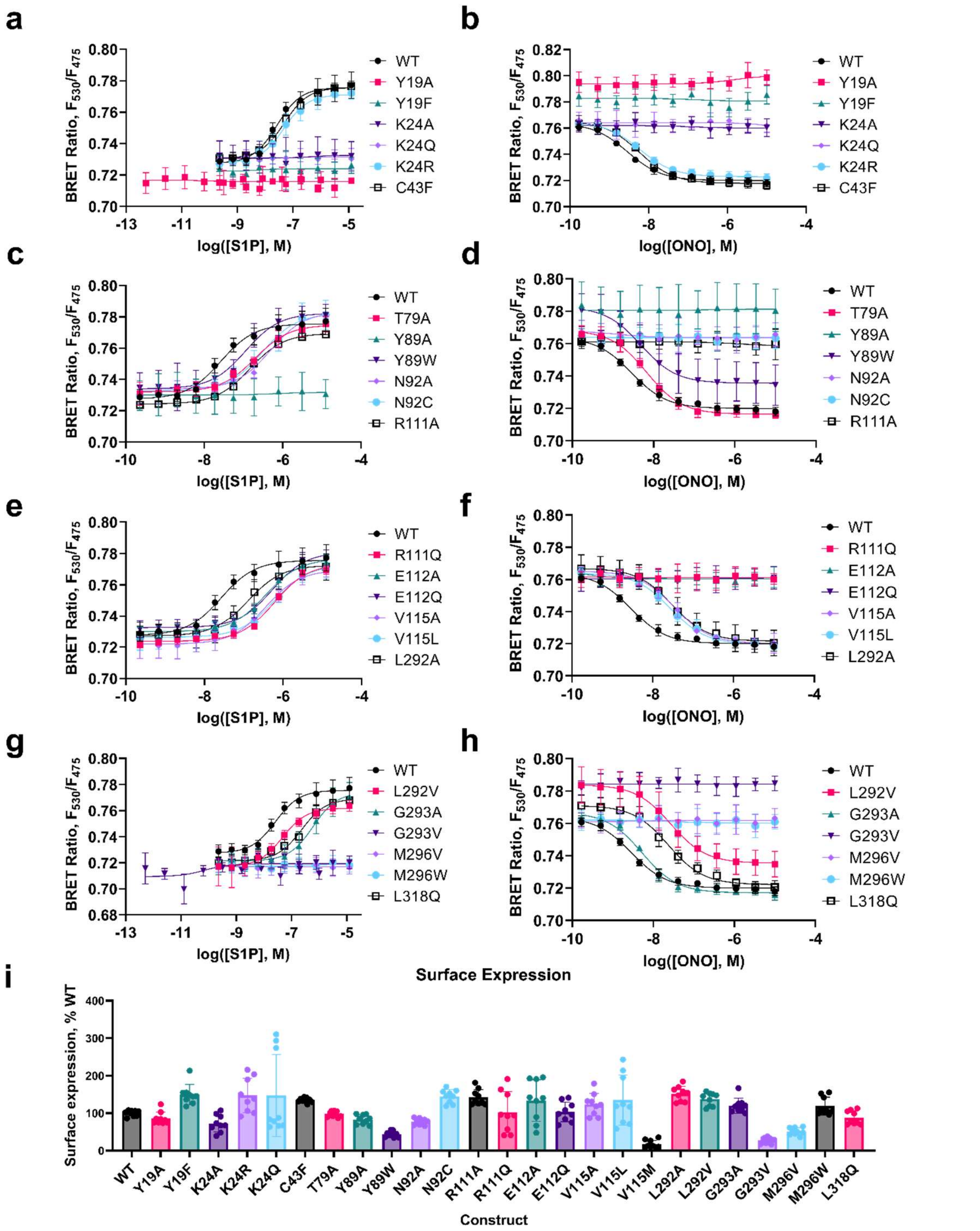


**Supplementary Fig. 6 Dose-response curves for S1P and ONO-5430608 and cell surface expression for WT and mutants of S1P$_5$. a,c,e,g** S1P- stimulated cAMP reduction at WT S1P$_5$ and mutants, measured by BRET-based EPAC sensor. **b,d,f,h** Inhibition of forskolin-stimulated cAMP reduction by ONO-5430608 at WT S1P$_5$ and mutants, measured by BRET-based EPAC sensor. Each data point represents mean±s.d. for 3 independent experiments performed in triplicate. **i** Cell surface expression of the HA-tagged WT S1P$_5$ and mutants as determined by ELISA. Bar heights represent mean ± s.d. for 3 independent experiments performed in triplicate (all data points are shown).

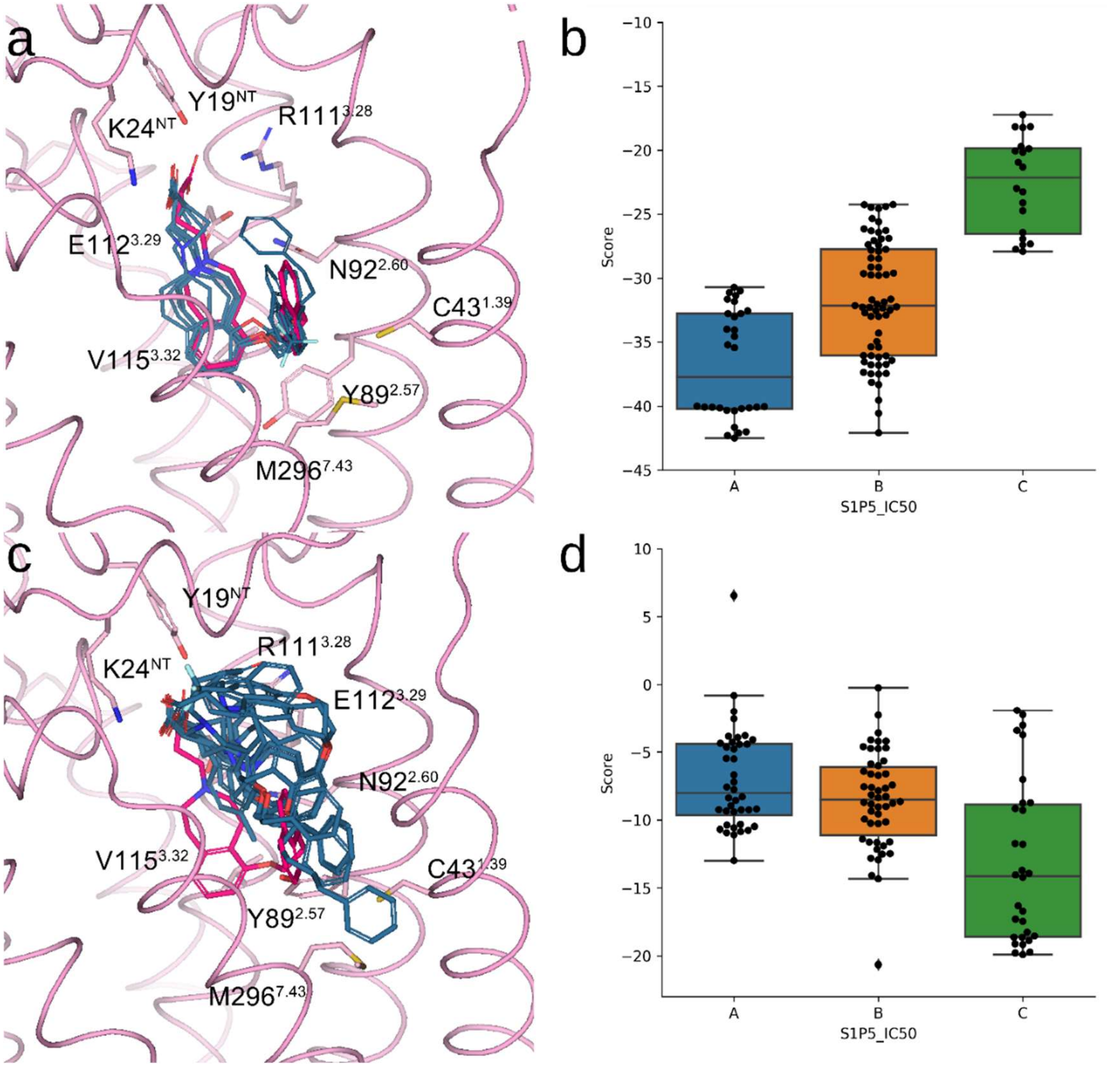


**Supplementary Fig. 7 Ligand docking simulations. a,c** Overlay of ligand binding poses (one highest-score pose per ligand) for all group 'A' ligands docked in the $S1P_5$ crystal structure (downward conformation of $Y89^{2.57}$) (**a**) or in a metaMD snapshot with an upward conformation of $Y89^{2.57}$ (**c**). **b,d** Clustering of docking scores for all tested ligands (5 trials per ligand) corresponding to docking runs described in (**a,c**), respectively. All ligands are grouped according to their $S1P_5$ affinity: 'A' (1 nM < $IC_{50}$ < 100 nM), 'B' (100 nM < $IC_{50}$ < 1 μM), 'C' (>1 uM).

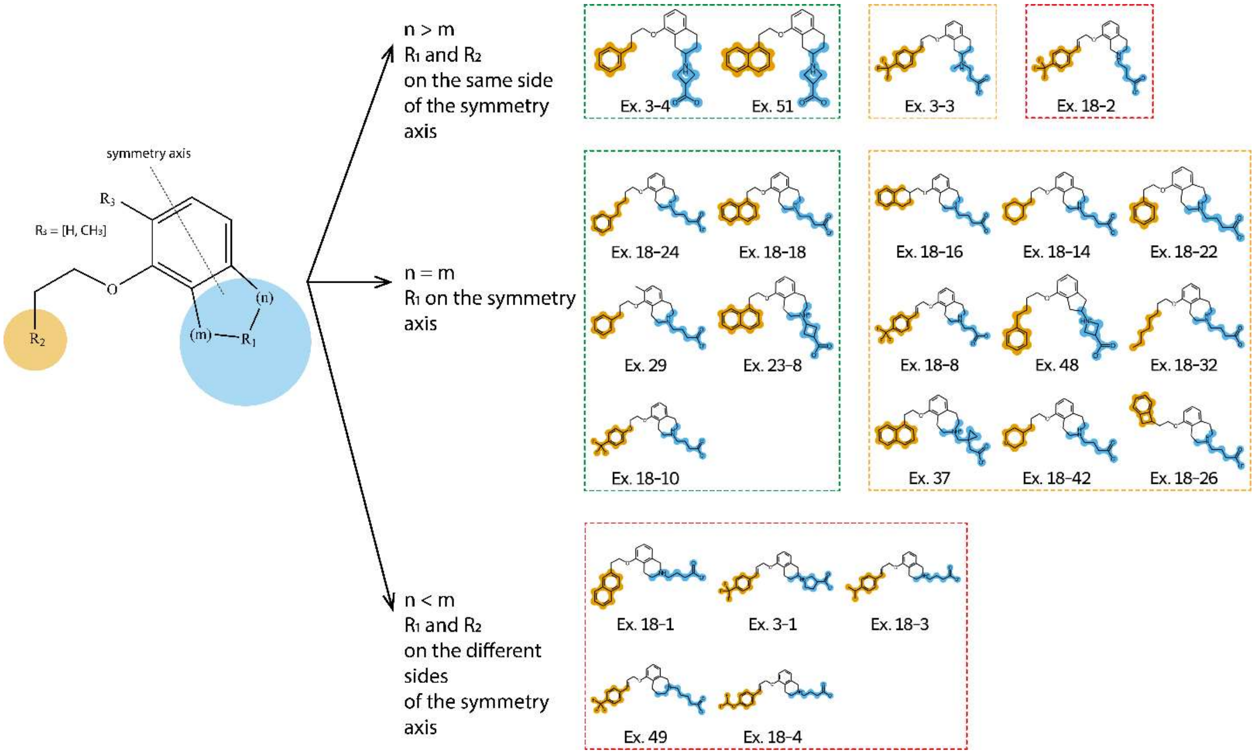


**Supplementary Fig. 8 Substituent decomposition analysis of ONO-5430608 ligand series**[1]. The double ring system symmetry axis is shown in dotted line. Rows represent different substituent symmetry about the double ring system symmetry axis: $R_1$ and $R_2$ are on the same side (top row), $R_1$ is on the axis (middle row), $R_1$ and $R_2$ are on the different sides (bottom row). Ligand groups are outlined with respect to their affinities: group 'C' ($IC_{50}$ between 1 and 3 μM) in red, group 'B' ($IC_{50}$ between 100 nM and 1 μM) in yellow, and group 'A' ($IC_{50}$ between 1 and 100 nM) in green.

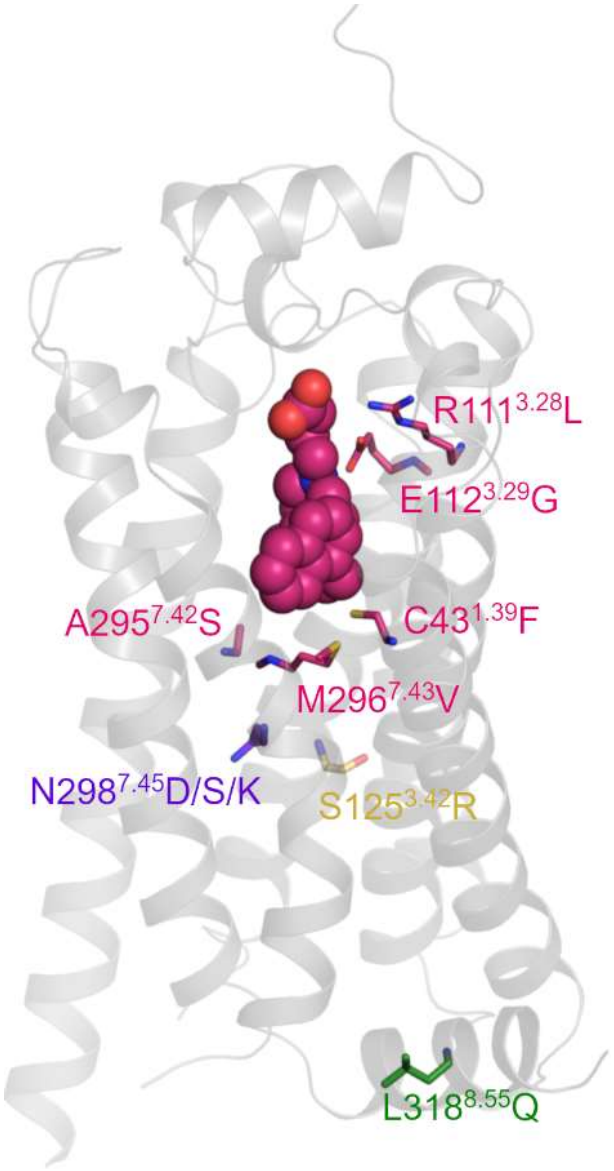


**Supplementary Fig. 9 Examples of naturally occurring missense SNVs in S1P$_5$, mapped on the receptor structure.** SNVs in the ligand binding pocket are shown in pink, in the sodium site - in purple blue, disrupting hydrogen bond - in yellow, disrupting $G_{12/13}$ signaling - in green.

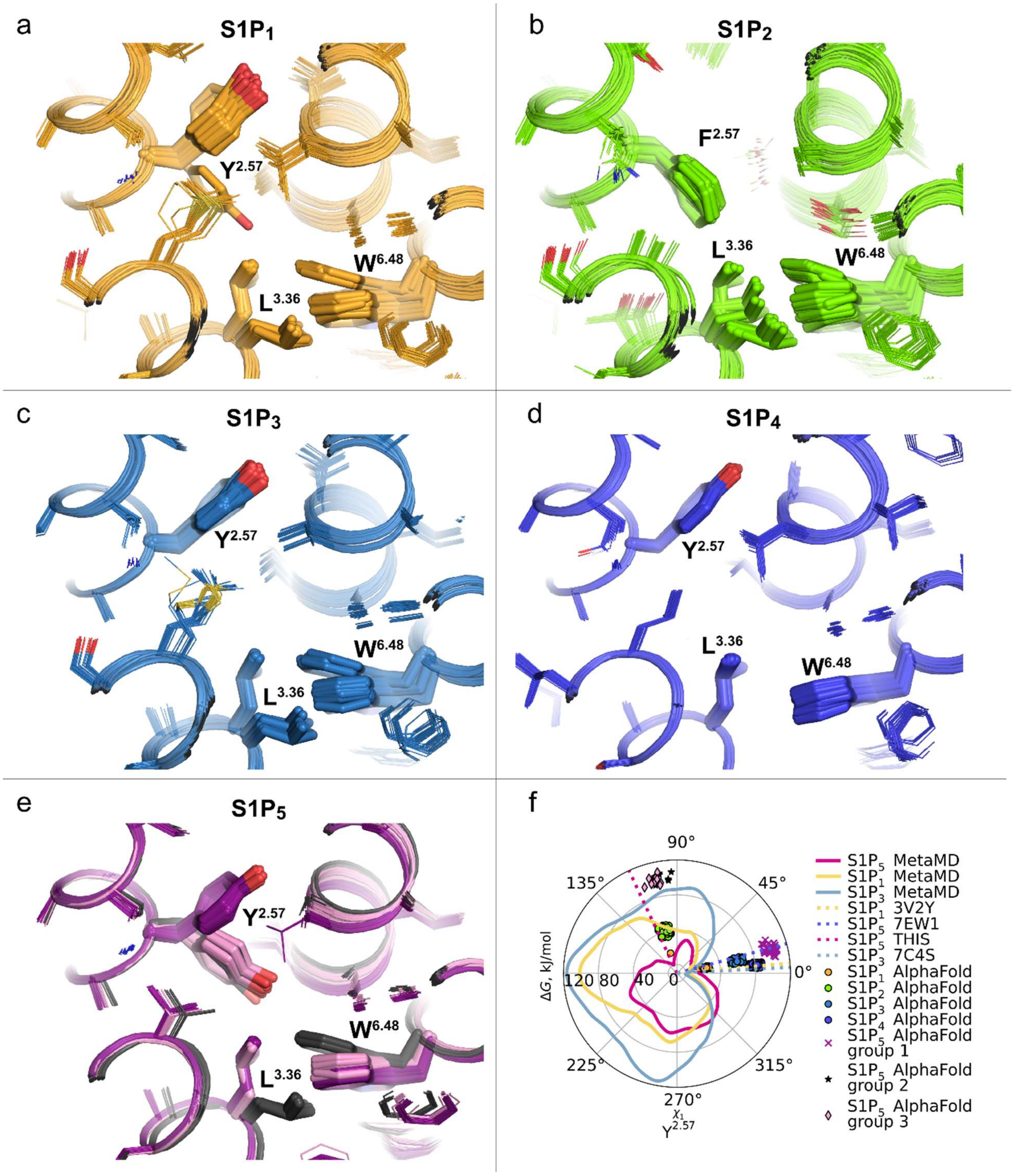


**Supplementary Fig. 10 AlphaFold prediction of S1PRs. a-e** Conformations of $Y^{2.57}$ and the dual toggle switch $L^{3.36}$-$W^{6.48}$ in 50 AlphaFold models of $S1P_{1-5}$ subtypes, respectively. Three distinct conformations of $S1P_5$ are shown in dark violet, light violet, and black. **f** Free energy profiles of the $Y^{2.57}$ side chain torsion angle $\chi_1$ in $S1P_1$ (yellow line), $S1P_3$ (blue line) and $S1P_5$ (pink line) calculated by metaMD. Dotted lines correspond to $Y^{2.57}$ conformations in experimental structures. Individual points correspond to $Y^{2.57}$ conformations in AlphaFold models.

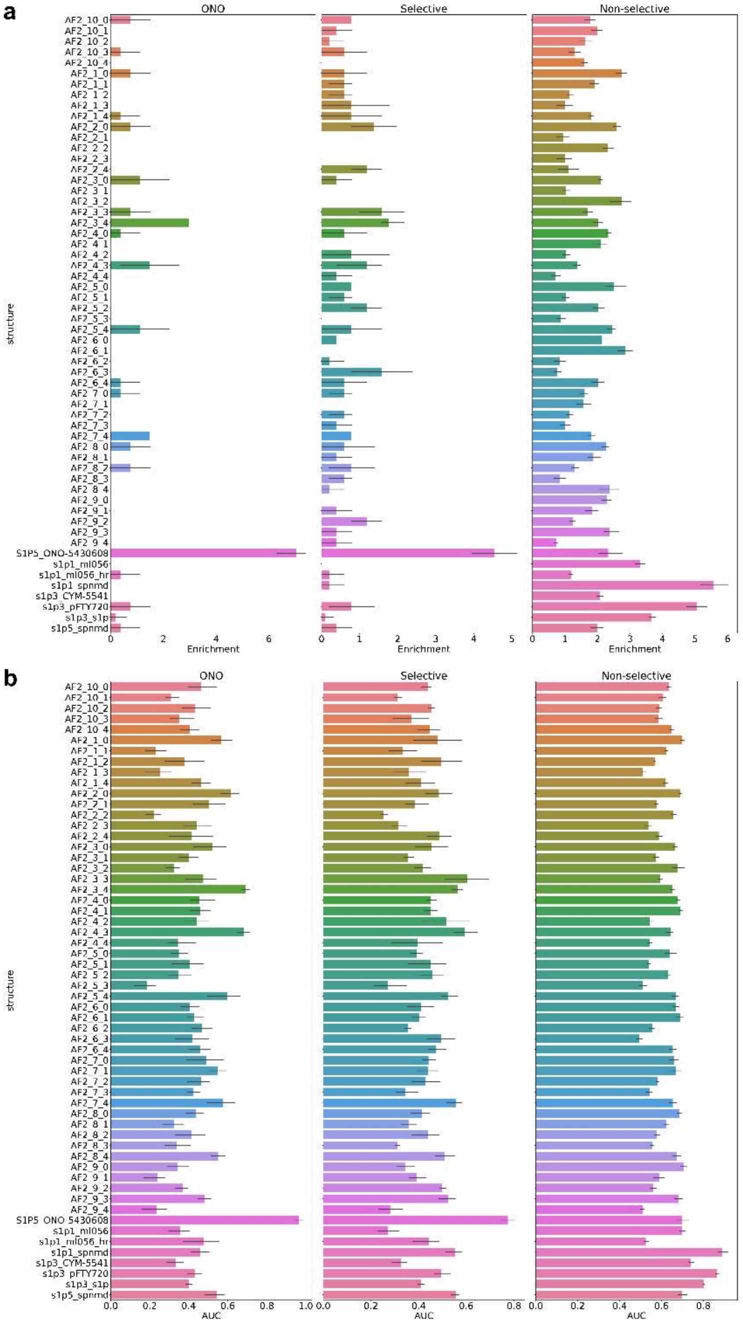


**Supplementary Fig. 11 Virtual ligand screening benchmark comparison of AlphaFold models and experimental structures**. **a** 10%-enrichment- score for 50 S1PR AlphaFold models and available experimental structures in three different benchmark sets. **b** ROC-AUC score for 50 S1PR AlphaFold models and available experimental structures in three different benchmark sets. Bar heights represent mean ± s.d. for 3 docking trials with effort=1. Benchmark sets from left to right: 'ONO' – ligands from ONO-5430608 series[1], 'Selective' – selective S1P$_5$ ligands[1,2], 'Non-selective' - S1PR ligands (from ChEMBL[3]). Experimental structures used for screening: S1P5_ONO-5430608 (this work, PDB ID 7YXA), s1p1_ml056 (PDB ID 3V2W), s1p1_ml056_hr (PDB ID 3V2Y), s1p1-spnmd (PDB ID 7EVY), s1p3_CYM-5541 (PDB ID 7EW4), s1p3_pFTY720 (PDB ID 7EW2), s1p3-s1p (PDB ID 7C4S), s1p5-spnmd (PDB ID 7EW1).

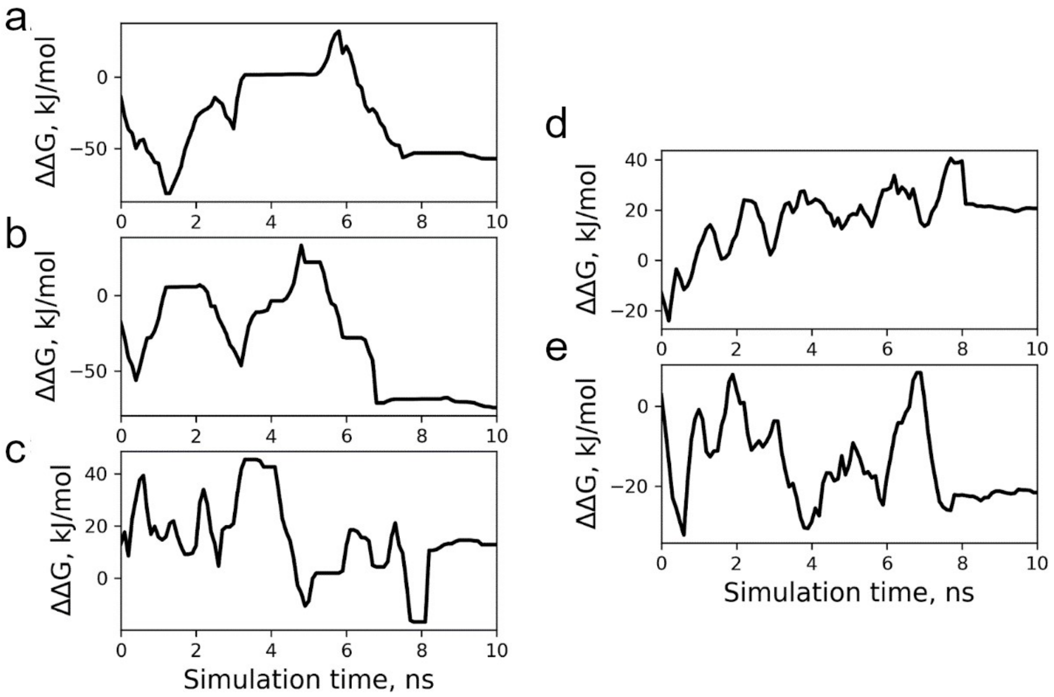


**Supplementary Fig. 12 Convergence of metaMD simulations. a-c** Convergence of metaMD free energy profiles of the $Y^{2.57}$ side chain torsion angle $\chi_1$ in $S1P_1$ (**a**), $S1P_3$ (**b**), and $S1P_5$ (**c**). **d,e** Convergence of metaMD free energy profiles of the $L^{3.36}$ side chain torsion angle $\chi_1$ in $S1P_5$ with upward (**d**) and downward (**e**) oriented $Y^{2.57}$. The convergence was estimated by tracking the difference between two distinct regions of the free energy profiles (ΔΔG) corresponding to the orientations of $Y^{2.57}$ in the $S1P_1$, $S1P_3$, and $S1P_5$ structures in (**a-c**) and to the orientations of $L^{3.36}$ in the active and inactive $S1P_5$ structures (**d,e**) as a function of simulation time. In case of convergence, this difference should not change with the progress of simulations as the systems diffuse freely along the reaction coordinate as observed during the last 2 ns of simulations on average.

### Supplementary Tables

**Supplementary Table 1 Crystallographic data collection and refinement statistics.**

| **Sample** | **$S1P_5$-ONO5430608** | |
|---|---|---|
| PDB ID | **7YXA** | |
| **Data collection** | | |
| Number of frames / crystals | 6,818 / 7,492 | |
| Space group | $P\,2_1\,2_1\,2_1$ | |
| Cell dimensions | | |
| *a*, *b*, *c* (Å) | 59.8, 103.4, 187.9 | |
| α, β, γ (°) | 90, 90, 90 | |
| Resolution (Å) | 30–2.2 (2.28–2.20) | |
| No. total reflections | 7,236,474 | |
| No. unique reflections | 60,645 | |
| $R_{split}$ (%) | 24.4 (71.8) | |
| $R_{meas}$ (%) | N/A | |
| Mean $I/\sigma I$ | 3.8 (1.5) | |
| Completeness (%) | 100.0 (100.0) | |
| Multiplicity | 119.3 (112.0) | |
| CC* (%) | 97.9 (51.6) | |
| **Refinement** | | |
| No. reflections / test set | 59,756 (4,564) | |
| $R_{work}/R_{free}$ (%) | 29.4 / 35.7 | |
| No. atoms | | |
| Protein | 2,804 | 2,808 |
| Ligand | 30 | 30 |
| Lipid and other[b] | 219 | |
| Wilson *B*-factors ($Å^2$) | 26.6 | |
| Overall mean *B*-factors ($Å^2$) | A | B |
| $S1P_5$ | 30.6 | 33.6 |
| BRIL | 44.9 | 53.3 |
| Ligand | 24.4 | 28.3 |
| Lipid and other[b] | 35.2 | |
| R.m.s. deviations | | |
| Bond lengths (Å) | 0.015 | |
| Bond angles (°) | 1.41 | |
| Ramachandran stats (%)[c] | | |
| Favored | 97.54 | |
| Allowed | 2.46 | |
| Outliers | 0 | |
| MolProbity score[c] | 1.36 | |

[a]Values in parentheses are for highest-resolution shell.

[b]NAG molecule bound to N-terminal helixes in both chains is accounted in "other".

[c]As defined by Molprobity[4]

**Supplementary Table 2 Potencies ($pEC_{50}$) of S1P and ONO-5430608 at $S1P_5$ WT and mutants measured by BRET-based cAMP assay and cell surface expression.**

| Mutation | ONO-5430608 $pEC_{50}$, M | S1P $pEC_{50}$, M | Surface Expression, %WT |
|---|---|---|---|
| WT | 8.54±0.04 | 7.58±0.07 | 100±8 |
| $Y19^{NT}A$ | N/R | N/R | 86±17 |
| $Y19^{NT}F$ | N/R | N/R | 150±30 |
| $K24^{NT}A$ | N/R | N/R | 70±20 |
| $K24^{NT}Q$ | N/R | N/R | 150±110 |
| $K24^{NT}R$ | 8.23±0.04 | 7.34±0.07 | 150±50 |
| $C43^{1.39}F$ | 8.24±0.03 | 7.32±0.03 | 135±6 |
| $T79^{2.47}A$ | 8.17±0.04 | 6.70±0.06 | 98±8 |
| $Y89^{2.57}A$ | N/R | N/R | 83±13 |
| $Y89^{2.57}W$ | 8.31±0.14 | 6.95±0.10 | 44±8 |
| $N92^{2.60}A$ | N/R | 6.31±0.04 | 78±8 |
| $N92^{2.60}C$ | N/R | 6.52±0.08 | 145±18 |
| $R111^{3.28}A$ | N/R | 6.84±0.03 | 140±20 |
| $R111^{3.28}Q$ | N/R | 6.23±0.03 | 100±50 |
| $E112^{3.29}A$ | N/R | 6.45±0.05 | 130±60 |
| $E112^{3.29}Q$ | N/R | 6.28±0.06 | 100±30 |
| $V115^{3.32}A$ | 7.46±0.04 | 6.45±0.09 | 120±30 |
| $V115^{3.32}L$ | 7.55±0.03 | 6.24±0.07 | 140±70 |
| $V115^{3.32}M$ | N/A | N/A | 17±10 |
| $L292^{7.39}A$ | 7.51±0.09 | 6.89±0.09 | 152±20 |
| $L292^{7.39}V$ | 7.57±0.10 | 7.33±0.10 | 138±18 |
| $G293^{7.40}V$ | 8.28±0.03 | 6.22±0.11 | 120±20 |
| $G293^{7.40}A$ | N/R | N/R | 28±8 |
| $M296^{7.43}V$ | N/R | N/R | 52±10 |
| $M296^{7.43}W$ | N/R | N/R | 120±20 |
| $L318^{8.55}Q$ | 7.63±0.06 | 6.64±0.06 | 88±18 |

Data represent mean±s.d. of 3 independent experiments performed in triplicate.

**Supplementary Table 3 DNA sequence of the S1P$_5$ crystallization construct.**

```
ATGAAGACCATCATCGCTCTGTCCTACATCTTCTGCCTCGTGTTCGCCgactacaaggacgatgacgatgctg
ggcgcgccatggaatcgggactcttgcgtccggctcctgtctctgaggtcatcgttctgcattacaactacactggaaaactgaggggtgcgaggt
accagcctggagctggattgagagctgacgcagtggtctgcctggcagtttgtgcgttcatcgtgctcgagaacttggctgtgctgctcgtcctggg
aaggcacccaagattccatgctccgatgttcttgctgctcggttcactcaccttgagtgatttgctggctggcgctgcctacgcagcgaacatcctctt
gtcgggaccactgaccctcaagttgtccccggctctgtggttcgccagagagggtggcgtcttcgttgctctgactgccagcgtcctctctctgctcg
caattgcgttggaacgctccctgacaatggcacgccgtggaccagcaccggtgtccagcagaggacgtacgctcgctatggctgcagcagcat
ggggagtctcattgctgctcggtttgctgccagctctgggatggaactgcctgggaagactcgacgcctgttccactgttctgccgctctacgctaag
gcctacgttctcttctgcgtgttggccttcgtcggcatcctcgctgccatttgcgcattgtacgcgaggatctactgtcaggtgagagcaaacgctgct
gatctggaagacaattgggaaactctgaacgacaatctcaaggtgatcgagaaggctgacaatgctgcacaagtcaaagacgctctgaccaa
gatgagggcagcagccctggacgctcagaaggccactccacctaagctcgaggacaagagcccagatagccctgaaatgaaagactttcg
gcatggattcgacattctggtgggacagattgatgatgcactcaagctggccaatgaagggaaagtcaaggaagcacaagcagccgctgagc
agctgaagaccacccggaatgcatacattcagaagtacctgcgccgtaaaccacgtagcctggctctcttgaggacactctctgttgtgctgctcg
ctttcgtcgcctgctggggaccactgttcttgctgctcttgctggacgtcgcatgccctgcgcgtacgtgtcccgttctcttgcaagccgatcctttcttgg
gtctggctatggccaactctctgctcaaccccatcatttacacattgacgaaccgcgacctgcgtcacgctttgctgaggctggtgggaagacctC
TGGAGGTGCTCTTCCAGGGTCCCCACCATCATCACCATCATCACCACCACCACTAA
```

Sequences of the cleaved tags and the stop codon are shown in capital letters.

**Supplementary Table 4 Optimization of CrystFEL parameters and data quality improvement.**

| | | A | B | C | D | E | F |
|---|---|---|---|---|---|---|---|
| **Peak search** | SNR | 4 | 2.7 | 4 | 4 | 4 | 4 |
| | Threshold | 100 | 30 | 30 | 30 | 30 | 30 |
| | median-filter | - | - | 5 | 5 | 5 | 5 |
| **Integration** | regime | rings-nograd | rings-nograd | rings-nograd | rings-grad | rings-grad | rings-grad |
| | int-radius | 4,5,8 | 4,5,8 | 4,5,8 | 4,5,8 | 4,5,8 | 3,5,8 |
| | local-bg-radius | - | - | - | - | 5 | 5 |
| **merging** | model | unity | unity | unity | unity | unity | unity |
| | pushres | 3 | 3 | 3 | 3 | 3 | 3 |
| | iterations | 1 | 1 | 1 | 1 | 1 | 1 |
| **Nframes** | | 1960 | 5007 | 6605 | 6605 | 6594 | 6600 |
| **Ncrystals** | | 2036 | 5275 | 7189 | 7185 | 7200 | 7205 |
| **Resolution** | low | 30-5.4 | 30-5.4 | 30-5.4 | 30-5.4 | 30-5.4 | 30-5.4 |
| | high | 2.59-2.50 | 2.59-2.50 | 2.59-2.50 | 2.59-2.50 | 2.59-2.50 | 2.59-2.50 |
| | overall | 30-2.50 | 30-2.50 | 30-2.50 | 30-2.50 | 30-2.50 | 30-2.50 |
| **Rsplit** | low | 42.34 | 33.74 | 29.71 | 19.44 | 19.12 | 20.55 |
| | high | 241.33 | 206.51 | 186.04 | 61.74 | 60.03 | 48.14 |
| | overall | 57.01 | 48.86 | 44.7 | 23.82 | 23.62 | 21.06 |
| **CC*** | low | 0.967 | 0.975 | 0.979 | 0.981 | 0.98 | 0.978 |
| | high | 0.554 | 0.587 | 0.618 | 0.666 | 0.676 | 0.716 |
| | overall | 0.964 | 0.974 | 0.978 | 0.984 | 0.983 | 0.983 |
| **Redundancy** | low | 96.8 | 314.1 | 503.8 | 149.8 | 148.9 | 124.9 |
| | high | 34 | 112 | 182.6 | 153.7 | 153.7 | 149.8 |
| | overall | 49.1 | 160.6 | 259.8 | 156.3 | 156 | 143.2 |
| **I/sigma** | low | 2.32 | 2.83 | 3.19 | 6.73 | 6.76 | 7.29 |
| | high | 0.47 | 0.52 | 0.58 | 1.87 | 1.88 | 2.41 |
| | overall | 1.38 | 1.61 | 1.78 | 4.12 | 4.12 | 4.97 |

**Supplementary Table 5 Primers used in this study**

| Primer Name | Forward Primer (5'-to-3') | Reverse Primer (5'-to-3') |
|---|---|---|
| $S1P_5$ crystallization construct | | |
| M1-V321 | GATGCTGGGCGCGCCATGGAATCG GGACTCTTGCG | GCACCTCCAGAGGTCTTCCCACCAG CCTCAGCAAAGCG |
| A223-Bril-R241 | GTCAGGTGAGAGCAAACGCTGCTG ATCTGGAAGACAATTGGG | GGCTACGTGGTTTACGGCGCAGGTA CTTCTGAATGTATGCATTCC |
| Functional tests primers | | |
| L292A | CTC CTG CAG GCC GAT CCC TTC GCC GGA CTG GCC ATG GCC AAC TC | GAG TTG GCC ATG GCC AGT CCG GCG AAG GGA TCG GCC TGC AGG AG |
| Y19A | CGAGGTCATCGTCCTGCATgccAAC TACACCGGCAAGCTCC | GGAGCTTGCCGGTGTAGTTggcATGC AGGACGATGACCTCG |
| Y19F | CGAGGTCATCGTCCTGCATttcAACT ACACCGGCAAGCTCC | GGAGCTTGCCGGTGTAGTTgaaATGC AGGACGATGACCTCG |
| K24A | CTGCATTACAACTACACCGGCgccCT CCGCGGTGCGCG | CGCGCACCGCGGAGggcGCCGGTGT AGTTGTAATGCAG |
| K24Q | CTGCATTACAACTACACCGGCcagC TCCGCGGTGCGCG | CGCGCACCGCGGAGctgGCCGGTGT AGTTGTAATGCAG |
| K24R | CTGCATTACAACTACACCGGCcggC TCCGCGGTGCGCG | CGCGCACCGCGGAGccgGCCGGTGT AGTTGTAATGCAG |
| C43F | GCCGACGCCGTGGTGttcCTGGCGG TGTGCGCC | GGCGCACACCGCCAGgaaCACCACG GCGTCGGC |
| M296W | CCTTCCTGGGACTGGCCtggGCCAA CTCACTTCTGAACCCC | GGGGTTCAGAAGTGAGTTGGCccaG GCCAGTCCCAGGAAGG |
| M296V | CCTTCCTGGGACTGGCCgtgGCCAA CTCACTTCTGAACCCC | GGGGTTCAGAAGTGAGTTGGCcacG GCCAGTCCCAGGAAGG |
| Y89A | GGCAGGCGCCGCCgccGCCGCCAA CATCCTACTGTC | GACAGTAGGATGTTGGCGGCggcGG CGGCGCCTGCC |
| Y89W | GGCAGGCGCCGCCtggGCCGCCAA CATCCTACTGTC | GACAGTAGGATGTTGGCGGCccaGG CGGCGCCTGCC |
| N92A | CCGCCTACGCCGCCgccATCCTACT GTCGGGGCCG | CGGCCCCGACAGTAGGATggcGGCG GCGTAGGCGG |

| | | |
|---|---|---|
| N92C | CCGCCTACGCCGCCtgcATCCTACT GTCGGGGCCG | CGGCCCCGACAGTAGGATgcaGGCG GCGTAGGCGG |
| E112A | CGCTCTGGTTCGCACGGgccGGAGG CGTCTTCGTGGC | GCCACGAAGACGCCTCCggcCCGTG CGAACCAGAGCG |
| E112Q | CGCTCTGGTTCGCACGGcagGGAG GCGTCTTCGTGGC | GCCACGAAGACGCCTCCctgCCGTGC GAACCAGAGCG |
| V115M | GCACGGGAGGGAGGCatgTTCGTGG CACTCACTGCG | CGCAGTGAGTGCCACGAAcatGCCTC CCTCCCGTGC |
| V115L | GCACGGGAGGGAGGCctgTTCGTGG CACTCACTGCG | CGCAGTGAGTGCCACGAAcagGCCT CCCTCCCGTGC |
| V115A | GCACGGGAGGGAGGCgccTTCGTG GCACTCACTGCG | CGCAGTGAGTGCCACGAAggcGCCT CCCTCCCGTGC |
| L318Q | TGCGCCACGCGCTCcagCGCCTGGT CTGCTGCG | CGCAGCAGACCAGGCGctgGAGCGC GTGGCGCA |
| R111Q | CGCGCTCTGGTTCGCAcagGAGGGA GGCGTCTTCGTG | CACGAAGACGCCTCCCTCctgTGCGA ACCAGAGCGCG |
| R111A | CGCGCTCTGGTTCGCAgccGAGGGA GGCGTCTTCGTG | CACGAAGACGCCTCCCTCggcTGCGA ACCAGAGCGCG |
| G293A | CAGGCCGATCCCTTCCTGgccCTGG CCATGGCCAACTC | GAGTTGGCCATGGCCAGggcCAGGA AGGGATCGGCCTG |
| G293V | CAGGCCGATCCCTTCCTGgtgCTGG CCATGGCCAACTC | GAGTTGGCCATGGCCAGcacCAGGA AGGGATCGGCCTG |
| L292V | CTGCAGGCCGATCCCTTCgtgGGAC TGGCCATGGCCA | TGGCCATGGCCAGTCCcacGAAGGG ATCGGCCTGCAG |

Mutated codons are shown in lowercase.
